# A Population-scale Single-cell Spatial Transcriptomic Atlas of the Human Cortex

**DOI:** 10.1101/2025.10.13.681959

**Authors:** Lei Han, Zhen Liu, Lifang Wang, Yuxuan Liu, Yanrong Wei, Jianyun Ma, Youzhe He, Yichen Zuo, Jiao Fang, Hui Yang, Xiang Zou, Zihan Wu, Min Wang, Wei Liu, Li Gao, Yuyang Liu, Xinxiang Song, Yao Santo Zhang, Junjie Lei, Huanhuan Li, Leiting Li, Yingjie An, Bufan Jin, Yanqing Zhong, Qinwen Chai, Quyuan Tao, Xing Tan, Youning Lin, Ruiyi Zhang, Shengpeng Wang, Mingkai Chang, Bolin Yang, Ming Chen, Lijuan Mi, Liyuan Zhuang, Nini Yuan, Chao Li, Tao Huang, Xin Li, Cirong Liu, Yidi Sun, Liang Chen, Longqi Liu, Xun Xu, Chengyu Li, Hui Guo, Hao Li, Muming Poo, Wen-Biao Gan, Jianhua Yao, Wen Yuan, Shiping Liu, Zhiming Shen, Ying Mao, Wu Wei

## Abstract

The genetic and spatial determinants of cell type diversity in human cerebral cortices remain poorly defined. Here, we present a population-level single-cell spatial transcriptomic atlas of human cortices from 71 donors across the lifespan. We identified 906 layer-specific genes showing conserved and divergent laminar expression patterns between humans and other species. Spatial analysis revealed neuronal vulnerability and glial activation during aging, together with a decline in the proportion of superficial SST neurons and their interactions with other cells. Disease-associated genes exhibited high cell-type and layer-specific expression, implicating the pathogenic role of spatially specific gene expression. Spatial cis-eQTL analysis identified regulatory variants linked to genes related to diseases like Tourette syndrome. Cross-species comparison demonstrated glial expansion in the human cortex, accompanied by enhanced neuron–glia communication via the neuregulin signaling. Together, we provide a comprehensive single-cell atlas of the human cortex that is essential for understanding aging, evolution, and disease pathogenesis.

## Introduction

The human cerebral cortex is characterized by an intricate laminar and columnar organization that serves as the neural substrate for higher cognitive functions ^1–3^. A comprehensive understanding of cortical architecture at the molecular level is crucial, as spatial patterns of gene expression underlie neural circuit organization and brain function ^4–7^. Despite the cortex’s six well-defined layers and numerous specialized regions, the precise spatial arrangement of its diverse cell types and gene expression programs remains incompletely characterized. Single-cell transcriptomic studies have revealed remarkable cellular diversity in the human cortex ^8–12^, yet these dissociative approaches inherently lose information about each cell’s location in situ. Fundamental questions thus persist about how specific cell types and genes are spatially distributed across cortical layers and regions in the intact human cortex.

Spatial transcriptomics technologies now enable profiling of gene expression within intact tissue sections while preserving cellular positional information. Early applications in the human cortex identified genes with layer-enriched expression and refined laminar markers in individual cortical areas. For example, spatial transcriptomic analyses of the dorsolateral prefrontal cortex (DLPFC) from small adult cohorts (3 donors ^13^ and 10 donors ^14^) using 10x Genomics Visium platform revealed distinct layer-specific gene signatures and highlighted clinically relevant expression patterns. However, these pioneering studies were limited in sample size and offered only coarse spatial resolution, far from single-cell resolution. Conversely, multiplexed error-robust fluorescence in situ hybridization (MERFISH) achieves single-cell resolution ^15–17^ but is restricted to preselected gene panels, limiting its ability to capture the full transcriptomic landscape.

Current spatial transcriptomic studies remain limited in scope, with restricted coverage of cortical regions and insufficient representation across the human lifespan. It remains unclear whether the laminar molecular architecture identified locally is conserved across the entire cortex or exhibits region-specific variation, and how the spatial organization of gene expression changes throughout life. Moreover, recent single-cell expression quantitative trait locus (eQTL) studies have revealed cell type-specific effects of genetic variants ^18,19^, yet lack integration with spatial transcriptomic data to resolve these effects in situ. Additionally, the recent release of spatial transcriptomic atlases for the macaque and mouse brains ^20–22^ underscores the urgency of establishing a comparable human atlas for systematic cross-species analysis. These considerations underscore the need for a spatial transcriptomic resource of human cortex that is both multi-regional and population-scale, with single-cell resolution and unbiased transcriptome-wide coverage. The recent development of Stereo-seq, a spatial barcoding-based platform that achieves single-cell resolution with whole-transcriptome coverage ^20,23,24^, offers a unique opportunity to systematically dissect the molecular architecture of the human cortex, linking anatomical structure to transcriptional function across spatial and temporal contexts with potential genetic and evolutionary regulation.

In this study, we constructed a population-scale spatial transcriptomic atlas of the human cerebral cortex at single-cell resolution. By integrating Stereo-seq, single-nucleus RNA sequencing (snRNA-seq) and whole-genome sequencing (WGS), we profiled all four major cortical lobes from 71 donors spanning the human lifespan. This multi-omics approach enabled the in situ annotation of over 3.4 million cells and the incorporation of individual genetic information. Using this resource, we mapped layer-enriched gene expression signatures across cortical lobes and identified spatial patterns modulated by age. Further transcriptome variation analysis and cis-eQTL mapping revealed cell type- and layer-specific transcriptomic variation and genetic regulatory effects associated with neurological disorders. Cross-species comparison highlighted a pronounced expansion of glial-associated regulation in the human cortex. Together, this study provides an unprecedented spatiotemporal transcriptomic framework for the human cortex, addressing critical gaps in our understanding of brain organization and offering a foundation for exploring how aging, genetic factors, and evolution shape the human cortex.

## Results

### Population-scale spatial transcriptomic profiling of the human cortex

We analyzed 71 human samples derived from histologically normal cortical tissues surgically removed for accessing the disease tissue (**STAR Methods**), using a multi-omics approach combining Stereo-seq (81 sections from 58 donors), snRNA-seq (114 libraries from 58 donors), and WGS (66 libraries from 66 donors) (**Figure 1A**). The 71 donors include males (32 donors) and females (39 donors) of varying ages in five groups (Childhood, 1–12 years; Adolescence, 13–20 years; Early adulthood, 21–40 years; Late adulthood, 41–60 years; Aged, ≥ 61 years). The tissues covered four cortical lobes, including frontal lobe (FL), temporal lobe (TL), parietal lobe (PL), and occipital lobe (OL). Initially, snRNA-seq data were filtered by quality control process and then clustered and annotated (**STAR Methods**). Through unsupervised clustering analysis of the expression profiles of 458,914 nuclei and annotation with canonical marker genes ^25,26^, we identified 3 major classes (glutamatergic, GABAergic, and non-neuronal cells) and 25 subclasses of cell types (**Figures 1B** and **S1A**), including 13 glutamatergic neuron subclasses, 7 GABAergic neuron subclasses and 5 non-neuronal cell subclasses. Cell subclasses were further divided into 248 cell clusters using an iterative clustering strategy (**STAR Methods**). A random forest model was used to test the robustness of clustering results on 248 cell clusters, which showed high prediction accuracy (**Figure S1B**). Furthermore, cell clusters (tentatively referred to as “cell subtypes”) in this study were integrated and compared with two previously published human cortex snRNA-seq datasets ^8,27^, showing a high degree of overlap and consistency **(Figures S1C** and **S1D**).

**Figure 1.**
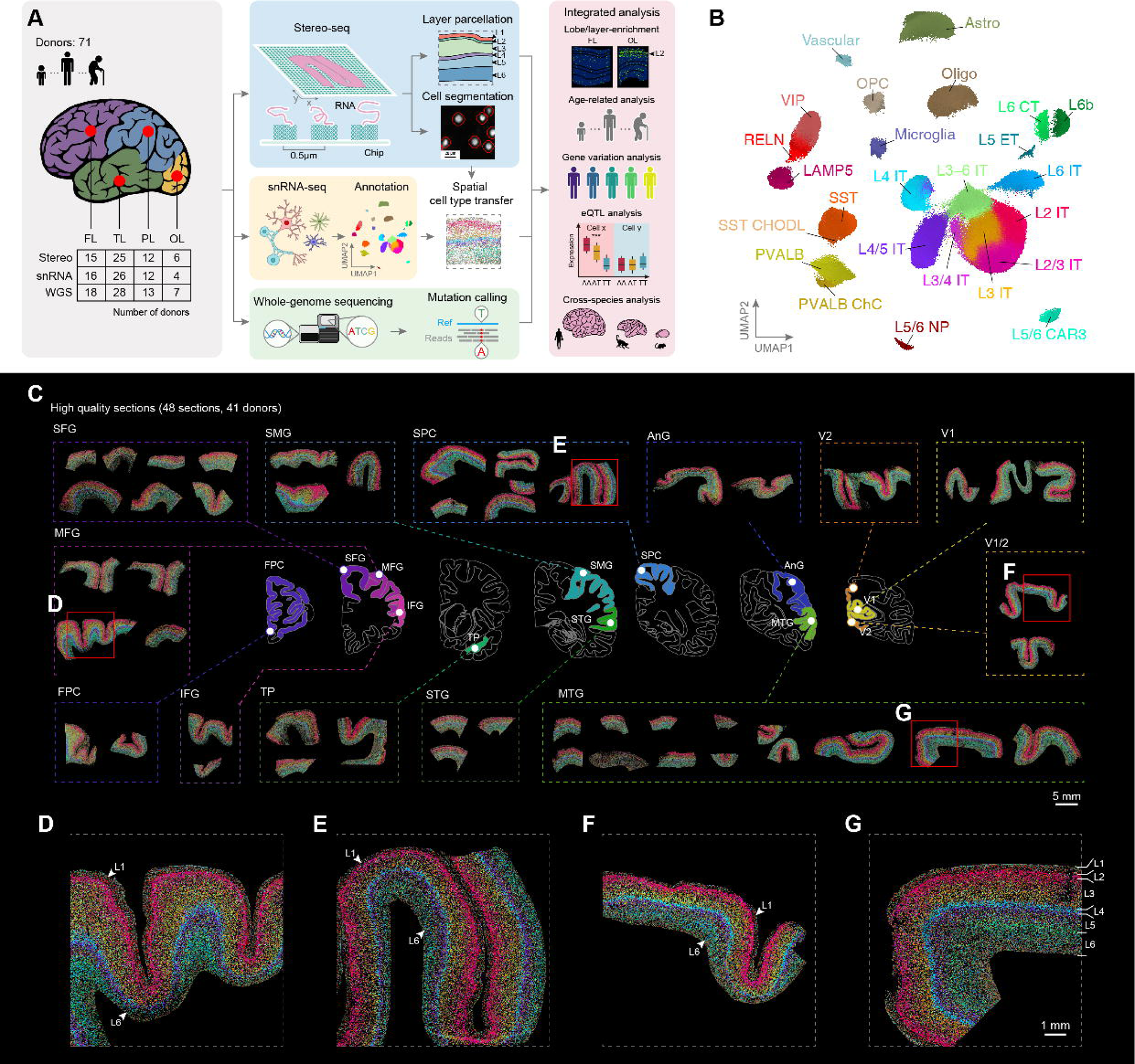
Multi-omics profiling of human cortical regions. (**A**) Schematic procedures showing the overview of sample collections and analysis of Stereo-seq, snRNA-seq and whole-genome sequencing. FL, frontal lobe; TL, temporal lobe; PL, parietal lobe; OL, occipital lobe. (**B**) Cell subclasses of snRNA-seq annotation in UMAP space. Cells were colored on 248 cell subtypes. (**C**–**G**) Spatial distribution of cells (colored by cell subtypes) on Stereo-seq sections. (**C**) Overview of 48 high-quality sections from 4 lobes, including 12 cortical regions. SFG, superior frontal gyrus; SMG, supramarginal gyrus; SPC, superior parietal cortex; AnG, angular gyrus; V1, primary visual cortex; V2, parastriate cortex; MFG, middle frontal gyrus; FPC, frontal polar cortex; IFG, inferior frontal gyrus; TP, temporal polar cortex; STG, superior temporal gyrus; MTG, middle temporal gyrus. Scale bar, 5 mm. (**D**–**G**) Zoomed-in views of regions labeled by red boxes in panel C. Scale bar, 1 mm. See also Figure S1.

To obtain single-cell resolution cell-type annotation for Stereo-seq data, we segmented single cells using watershed algorithm based on the nucleic-acid staining as described previously ^23^. This process yielded 3,415,566 segmented cortical cells across 81 sections, with an average of 970 unique molecular identifiers (UMIs) and 533 detected genes per cell. In total, this dataset captured expression of 34,949 genes (UMIs ≥ 2 in at least one cell, covering 94.5% protein coding and 90.4% lncRNA genes). Subsequently, the cell subtype annotations derived from the snRNA-seq data were transferred onto the segmented single cells using Spatial-ID, a graph convolution network (GCN)-based method ^28^ (**STAR Methods**). Additionally, we demonstrated that these spatial annotations showed high transcriptomic concordance with the snRNA-seq data (**Figures S1E** and **S1F**; **STAR Methods**), confirming the high accuracy of cell-type transfer. We visualized the spatial distribution of annotated cells in 48 high-quality sections (**Figures 1C**–**1G**; **STAR Methods**), revealing the laminar arrangement of cell subtypes across the human cortex. Overall, glutamatergic neurons demonstrated well-defined layer-enriched patterns, while GABAergic neurons and non-neuronal cells showed relatively dispersed distributions, consistent with previous observations in macaque and mouse cortices (**Figures S1G** and **S1H**). Nevertheless, several GABAergic and non-neuronal subtypes demonstrated distinct layer-(L)-enriched distributions. For example, among the somatostatin-positive (*SST*^+^) interneurons, various subtypes were preferentially enriched in L2/3, L4 and L5/6 respectively (**Figure S1I**), illustrating the fine-scale laminar heterogeneity in cortical cell subtypes.

### Layer-specific gene expression and cell-type composition across cortices

Cortical layer-specific gene expression patterns are fundamental to the cytoarchitectonic organization and functional specialization of distinct brain regions. However, previous studies for identifying genes with cortical layer-enriched expression (GLEEs) in human cortical tissues were limited by the sample size and spatial coverage, typically focusing on one or two cortical lobes ^13,29,30^. It remains unclear whether these identified GLEEs were conserved across all cortical lobes. Leveraging the generated Stereo-seq data, we performed a systematic characterization of GLEEs across four human cortical lobes to assess whether there were cortical region-dependent GLEEs in human cortex.

Overall, we identified 906 GLEEs across four cortical lobes, including 419 lobe-conserved GLEEs (C-GLEEs), 406 lobe-specific GLEEs (S-GLEEs) with restricted laminar expression in several specific lobes, and 81 GLEEs with different layer specificity in various lobes (**Figures 2A**–**2C**; **STAR Methods**). The majority of GLEEs reported in the previous single-lobe study ^29^ were recovered in our dataset, confirming the robustness of our approach. Additionally, applying the same method to 10x Visium data ^13^ from the DLPFC yielded GLEEs highly concordant with those we identified in the frontal lobe using Stereo-seq (**Figures S2A** and **S2B**). The GO analysis revealed distinct functional pathways among the three GLEE categories. C-GLEEs were enriched for corticogenesis-related processes (e.g., neuron projection development and trans-synaptic signaling), S-GLEEs for electrophysiological functions (e.g., ion channel activity), and the other complex GLEEs for structural connectivity (e.g., cell–cell adhesion and axonogenesis) (**Figure S2C**).

**Figure 2.**
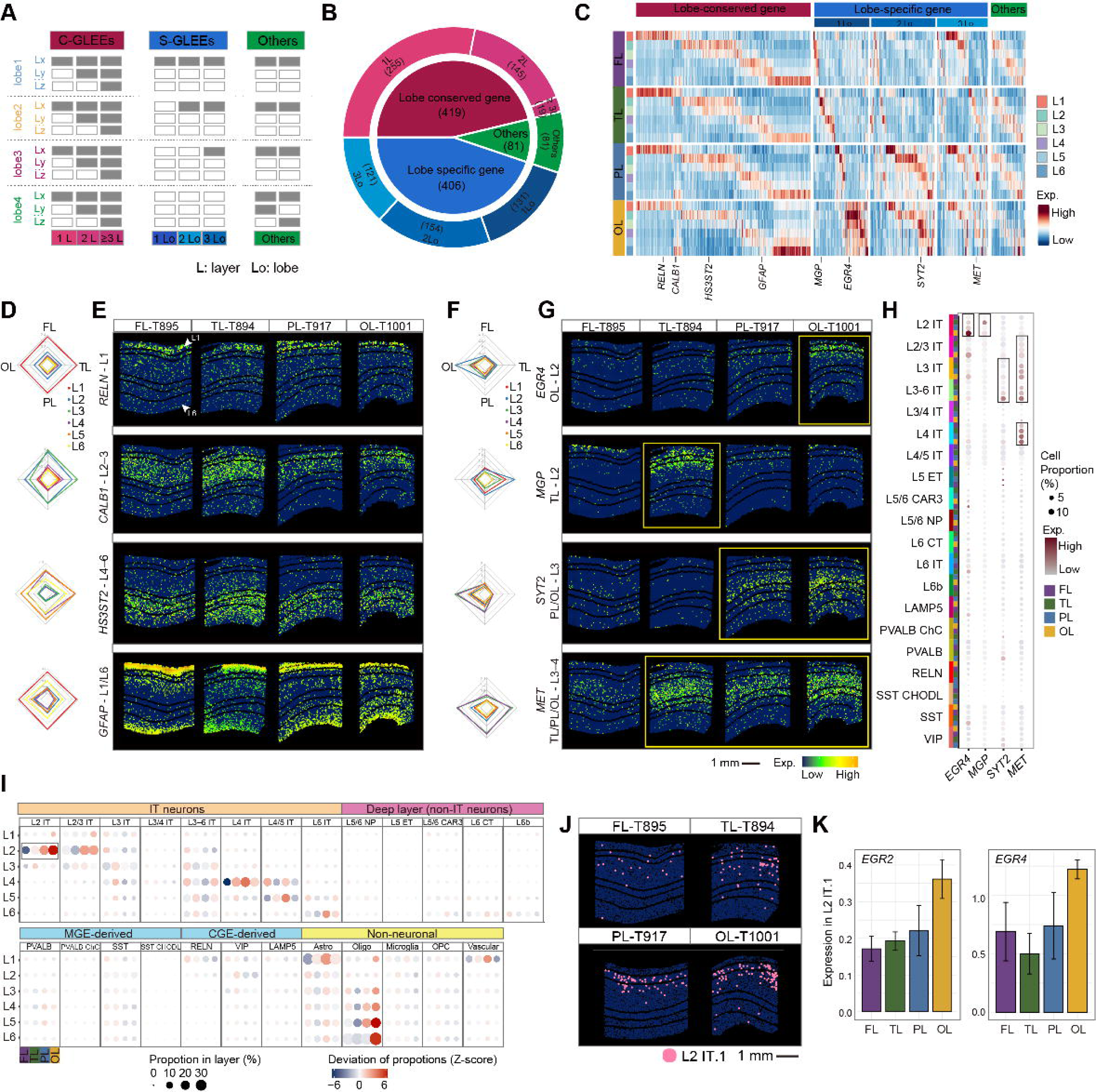
The layer distribution diversity of genes and cells across four cortical lobes. (**A**) Schematic diagram of 906 layer-enriched expressed genes (GLEEs) classification. GLEEs are categorized into three groups based on their cortical layer-enrichment patterns across the four cerebral lobes: (1) Lobe-conserved GLEEs (C-GLEEs), showing conserved laminar expression across all four cortical lobes, further subdivided into genes enriched in a single layer (1L-C-GLEEs), two layers (2L-C-GLEEs), or three or more layers (≥ 3L-C-GLEEs); (2) Lobe-specific GLEEs (S-GLEEs), genes enriched in specific subsets of lobes, further subdivided into single-lobe-specific (1Lo-S-GLEEs), two-lobe-specific (2Lo-S-GLEEs) and three-lobe-specific (3Lo-S-GLEEs); (3) Others: genes exhibiting complex expression patterns, present in all four lobes but with inconsistent layer-enrichment profiles. (**B**) Sunburst plot showing the categories of 906 GLEEs and the number of genes in each category. (C) Heatmaps show the mean expression of the top 50% of the sections in each lobe for the 906 GLEEs. (**D**–**G**) Spatial distribution of representative C-GLEEs and S-GLEEs. Radar chart showing the average expression of representative genes for C-GLEEs (**D**) and S-GLEEs (**F**) across each layer in all four cortical lobes. Spatial expression pattern of representative C-GLEEs (**E**) and S-GLEEs (**G**). Scale bar, 1 mm. (**H**) Dot plot showing the expression patterns of S-GLEEs from panel F in the snRNA-seq data. (**I**) Dot plot showing the differences between the proportion of cell subclasses across cortical layers and lobes. For each subclass, the proportion in a given layer and lobe was calculated as the number of cells of that subclass divided by the total number of cells in the same layer and lobe. Dot color represents the deviation (Z-score) from the average proportion across lobes (red: higher, blue: lower), and dot size indicates the average proportion. (**J**) Spatial distribution of L2 IT.1 neurons in representative sections of four cortical lobes. Scale bar, 1 mm. (**K**) Bar plot showing the mean expression levels of *EGR2* and *EGR4* within L2 IT.1 neurons across cortical lobes (mean ± SE). See also Figure S2.

Within the C-GLEEs, 255 genes (60.9%) displayed enrichment in a single cortical layer (1L-C-GLEEs) (**Figure 2B**). For example, *RELN*, a canonical marker for layer 1 (L1) neurons, was consistently expressed in L1 across all lobes (**Figures 2D**, **2E**, **S2D** and **S2E**), in agreement with its previously reported specificity in the DLPFC ^13^. Additionally, 145 (34.6%) genes were enriched in two layers (2L-C-GLEEs), such as *CALB1*, which was highly expressed in L2/3 across four lobes (**Figures 2D** and **2E**), and previously reported as a L2/3 specific gene in the occipital lobe (OL) ^29^. There was a smaller subset of C-GLEEs (19, 4.5%) enriched in three or more layers (≥ 3L-C-GLEEs), including *HS3ST2*, which was highly expressed in L4/5/6 across all four lobes (**Figures 2D** and **2E**). Intriguingly, we identified 26 C-GLEEs exhibiting enrichment in non-adjacent cortical layers. A notable example is *GFAP*, which displayed enriched expression patterns in both L1 and L6 (**Figures 2D** and **2E**).

Among the 406 S-GLEEs, 131 were uniquely enriched in a single cortical lobe (1Lo-S-GLEEs), including 13 in the frontal lobe (FL-S-GLEEs), 38 in the temporal lobe (TL-S-GLEEs), 29 in the parietal lobe (PL-S-GLEEs), and 51 in the occipital lobe (OL-S-GLEEs) (**Figure 2B**). For instance, *EGR2* and *EGR4*, two early-response transcription factors, were specifically expressed in L2 of the OL (**Figures 2F**, **2G**, **S2E**–**S2G**), consistent with their previously reported upregulation in the visual cortex in response to light stimulation ^31,32^. Similarly, *MGP* and non-coding gene *AC091230.1*, identified as TL-S-GLEEs, were specifically enriched in L2 and L6 of the TL, respectively (**Figures 2F**, **2G**, **S2F** and **S2G**). In addition, there were 154 S-GLEEs highly expressed in two cortical lobes (2Lo-S-GLEEs). For example, *SYT2* was highly expressed in L3 of the PL and OL. *SYT2* encodes a synaptic vesicle membrane protein involved in fast calcium dependent neurotransmitter release, and its enrichment in the visual cortex have been reported in prior studies ^29,33–36^ (**Figures 2F** and **2G**). Furthermore, there were 121 S-GLEEs highly expressed in three lobes (3Lo-S-GLEEs), such as *MET*, which showed specific expression in L3 and L4 across the TL, PL, and OL (**Figures 2F** and **2G**).

The other GLEEs showed more complex and heterogeneous laminar expression profiles across the four lobes. While broadly expressed across lobes, they lacked consistent layer specificity. For example, *PCP4* was enriched in L4–5 across four lobes but also showed strong expression in L2 of the TL and PL, deviating from prior reports describing it as L5-specific in the FL ^29^ and OL ^13^ (**Figure S2H**).

To investigate the cellular context of S-GLEEs, we analyzed both snRNA-seq and Stereo-seq data, which revealed that several S-GLEEs exhibit specific expression in glutamatergic neurons localized to their respective enriched cortical layers and lobes. For example, *EGR2* and *EGR4* was highly expressed in L2 intratelencephalic (IT) neurons and L2/3 IT neurons of the OL (**Figures 2H**, **S2I** and **S2J**), supporting its laminar and regional specificity. We next explored whether differences in S-GLEE expression across lobes are related to underlying variation in cell type composition. Along the rostro–caudal (R–C) axis, we observed a progressive increase in the ratio of glutamatergic (excitatory) to GABAergic (inhibitory) neuron (E:I ratio) across lobes (**Figure S2K**). A similar trend was observed with increasing cortical depth, consistent with previous research reports ^16^ (**Figure S2L**). Comparative analysis of cell-type distributions revealed that IT-neurons exhibit greater inter-lobe variability than non-IT excitatory neurons, inhibitory neurons, or non-neuronal cells ^16^ (**Figure 2I**). Notably, L2 IT neurons showed a gradual increase in abundance along the R-C axis, contributing to a lobe-specific increase in the E:I ratio within L2 (**Figures 2I**, **S2M** and **S2N**). Given their abundance and regional variation, we focused on L2 IT neurons and their association with S-GLEE. Notably, a lobe-specific L2 IT neurons subtype (L2 IT.1), specifically enriched in the OL (**Figure 2J**), was robustly detected in both snRNA-seq and Stereo-seq datasets (**Figure S2O**). This subtype exhibited strong expression of both *EGR2* and *EGR4*, suggesting a potential link between subtype-specific abundance and S-GLEE patterns in the OL (**Figures 2K** and **S2P**).

### Spatial diversity of age-related signatures in the human cerebral cortex

Our dataset spans a wide-range of ages from 1 to 76 years, allowing for a systematic investigation of the spatial distribution and cell-type specificity of age-related expression of genes (ARGs) in the human cerebral cortex (**Figure S3A**). To achieve this, we performed age-fitting analyses ^37^ (**STAR Methods**) on gene expression within each cortical layer, identified genes exhibiting significantly positive or negative correlation with the age. In total, we identified 1,814 ARGs strongly correlated with age (R ≥ 0.6), comprising 1,103 upregulated and 711 downregulated genes (**Figures 3A** and **3B**). These ARGs effectively distinguished the samples by its age, but not by the cortical lobe or gender (**Figures 3C**, **S3B** and **S3C**), indicating that age is the primary factor of their expression patterns.

**Figure 3.**
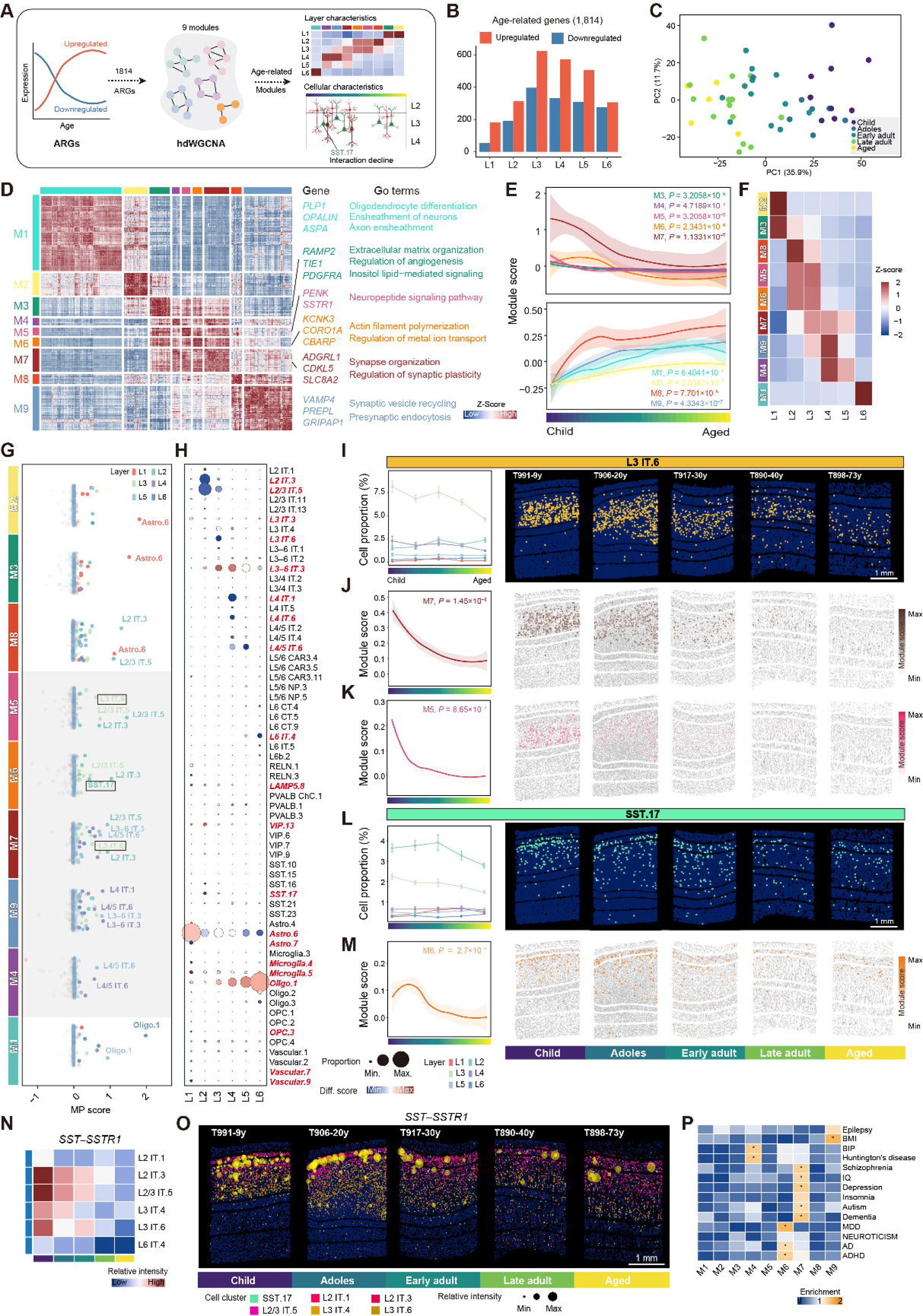
Spatial molecular and cellular characteristics of age-related gene (ARG) modules. (**A**) Schematic overview of the analytical workflow used to identify age-related signatures from Stereo-seq data. (**B**) Bar plot showing the number of ARGs in each cortical layer. (**C**) Principal component analysis (PCA) plot showing the distribution of donor Stereo-seq sections colored by different aging stages based on chronological age: childhood (Child, 1–12 years), adolescence (Adoles: 13–20 years), early adulthood (Early adult: 21–40 years), late adulthood (Late adult: 41–60 years), and aged (Aged, ≥ 61 years). PCA was performed using the expression profiles of all ARGs. (**D**) Heatmap of ARGs classified into 9 modules by hdWGCNA, each correlated with distinct GO biological processes. (**E**) Line chart depicting the module score (expression of module eigengenes) in each module across different aging stages. (**F**) Heatmap showing the scaled (Z-score) module score in each module across cortical layers. (**G**) Scatter plot depicting the Module Proportion Integrated Score (MP score; **STAR Methods**) of cell subtypes for each module within each layer. (**H**) Dot plot showing the proportion changes of cell subtypes in each cortical layer across aging stages. Dot size indicates the average proportion of cell subtypes in each cortical layer of all aging stages. Dot color reflects direction and magnitude of change with age (red, increase; blue, decrease; Diff. score, difference score). (**I**) Line chart depicting the layer-specific proportion dynamics of L3 IT.6 neurons across aging stages, alongside spatial visualization of L3 IT.6 neurons distributions. Scale bar, 1 mm. (**J** and **K**) Line chart showing the module scores of modules M5 and M7 (associated with L3 IT.6 neurons) across aging stages, alongside spatial visualization of M5 and M7 score distributions. (**L**) Line chart depicting the layer-specific proportion dynamics of SST.17 neurons across aging stages, alongside spatial visualization of SST.17 neurons distributions. Scale bar, 1 mm. (**M**) Line chart showing the module scores from corresponding modules (M6) related to SST.17 neurons across aging stages, alongside spatial visualization of M6 score distributions. (**N**) Heatmap showing cell–cell interaction intensity of the *SST*–*SSTR1* ligand–receptor pair across cell subtypes and aging stages. (**O**) Spatial visualization showing cell–cell interaction intensity of the *SST*–*SSTR1* ligand–receptor pair across aging stages. Yellow circle size reflects the interaction strength. Scale bar, 1 mm. (**P**) Heatmap depicting LDSC enrichment of GWAS traits and disorders related to brain health in ARG modules. *P*-values were derived from LDSC enrichment tests, and FDR-corrected *P*-values are overlaid on the heatmap (* for FDR < 0.05). See also Figure S3.

To further categorized these ARGs, we performed co-expression network analysis ^38^, grouping them into 9 distinct modules (M1–M9) based on their correlations with age (**STAR Methods**). These modules exhibited layer-specific expression patterns and were associated with distinct biological functions. For instance, ARGs in module M1 were prominently expressed in L6 and associated with myelination. ARGs in modules M2 and M3 showed high expression level in L1, enriched in processes such as angiogenesis and extracellular matrix organization, respectively. Meanwhile, ARGs in modules M5, M6 and M8 were more highly expressed in L2/3, with enriched functions related to stress-activated MAPK cascade, neuropeptide signaling, and cation transmembrane transport (**Figures 3D**–**3F**).

To determine whether ARGs within each module exhibit both layer and cell-type specificity, we evaluated the expression of ARGs in each module across various cell subtypes and cortical layers. Our analysis revealed that there were 5 modules (M4, M5, M6, M7 and M9) closely associated with specific neuronal subtypes, thus defined as neuron-associated modules (**Figure S3D**). For example, ARGs in module M7, which are involved in synapse organization and axon extension, were highly expressed in IT neurons (**Figure 3G**). Furthermore, we found a correlation between the expression patterns of ARGs in distinct module with the changes in cell composition. Notably, 4 neuron-associated modules (M4, M5, M6 and M7) generally showed downregulation with age, accompanied by a reduction in the proportion of specific neuronal subtypes (**Figures 3E** and **3G**). Among 20 subtypes with most pronounced age-dependent changes in their proportions, there were 9 glutamatergic and 3 GABAergic neuronal subtypes (**Figures 3H**, **S3E** and **S3F**). In particular, we found a significant age-related reduction in the proportion of L3 IT.6 subtype, indicating its age-dependent vulnerability (**Figure 3I**). Besides, we also observed age-dependent down-regulation of ARGs related to neuropeptide signaling (module M5) and synapse organization/axon extension (module M7) (**Figures 3J** and **3K**), indicating age-dependent decline in these two biological functions. The GO analysis also showed that this L3 IT.6 subtype expressed marker genes relating to these signaling pathways (**Figure S3G**). Compared with neuron-associated modules (M4, M5, M6, M7 and M9), we found that M1, M2 and M3 were highly associated with glia cells. For instance, ARGs in M1 and M2 were highly expressed in astrocytes and oligodendrocytes, respectively (**Figure 3G**) and were functionally enriched in glia cell proliferation and regulation of angiogenesis. Moreover, the age-dependent up-regulation of ARGs in these glia-associated modules was in line with the increasing proportion of glia cells in the cortex with age (**Figures 3H**, **S3E** and **S3F**).

Previous studies have found that degeneration of *SST*^+^ neurons are strongly associated with neurodegenerative diseases ^39–41^. Our study further identified a subtype of *SST*^+^ neurons (SST.17) that exhibited a pronounced reduction of their proportion with age. Intriguingly, ARGs involved in ion transport functions (module M6) were highly expressed in SST.17 neurons and showed parallel downregulation with the decline of SST.17 subtype during aging (**Figures 3L** and **3M**). Further analysis of cell–cell interaction based on ligand–receptor paired expression (**STAR Methods**) revealed that SST.17 neurons were physically proximal to glutamatergic neuron subtypes in L2/3, including L2 IT.3, L2/3 IT.5, and L3 IT.4 (**Figure 3N**). Notably, both the strength and number of these interactions decreased with age, especially for ligand–receptor pairs of *SST* and somatostatin receptors (*SSTR1*, *SSTR2* and *SSTR3*) (**Figures 3N**, **3O**, **S3H** and **S3I**). Among the three receptors, *SSTR1* showed the most prominent age-dependent down-regulation (**Figure S3J**), suggesting a reduction of synaptic interaction between SST.17 and L2/3 IT neurons in the aging brain. To investigate the relationship between various age-dependent gene modules and the brain health, we performed linkage disequilibrium score regression (LDSC) analysis (**STAR Methods**) using genome-wide association study (GWAS) data for brain health ^42^. Specifically, the module M6 showed significant enrichment (false discovery rate, FDR < 0.05) for Alzheimer’s disease (AD), attention deficit hyperactivity disorder (ADHD) and major depressive disorder (MDD) (**Figure 3P**), in line with the notion that the age-dependent reduction of SST.17 neurons may underlie neurological disorders.

### Contributions of cell type and layer location to spatial variation of gene expression

Our single-cell spatial transcriptomic atlas across a population-scale cohort offers a unique opportunity to study the sources of expression variation in the brain, especially cell locations. Thus we employed a linear mixed model (LMM) framework ^43^ to analyze the Stereo-seq sections of cortical tissues from various donors, quantitatively attributing gene expression variation to biological and technical factors, including the donor identity, cell type, cortical layer and region, as well as the age, gender and others (**Figure 4A**; **STAR Methods**). The percentage of the variance attributed to a given factor represents the factor’s contribution to the variation in gene expression. To validate the consistency of our results, we subsequently applied the LLM to snRNA-seq data. The percentages of contributing factors (non-residual factors) found for snRNA-seq and Stereo-seq data were highly correlated (**Figure S4A**), indicating high concordance between these datasets.

**Figure 4.**
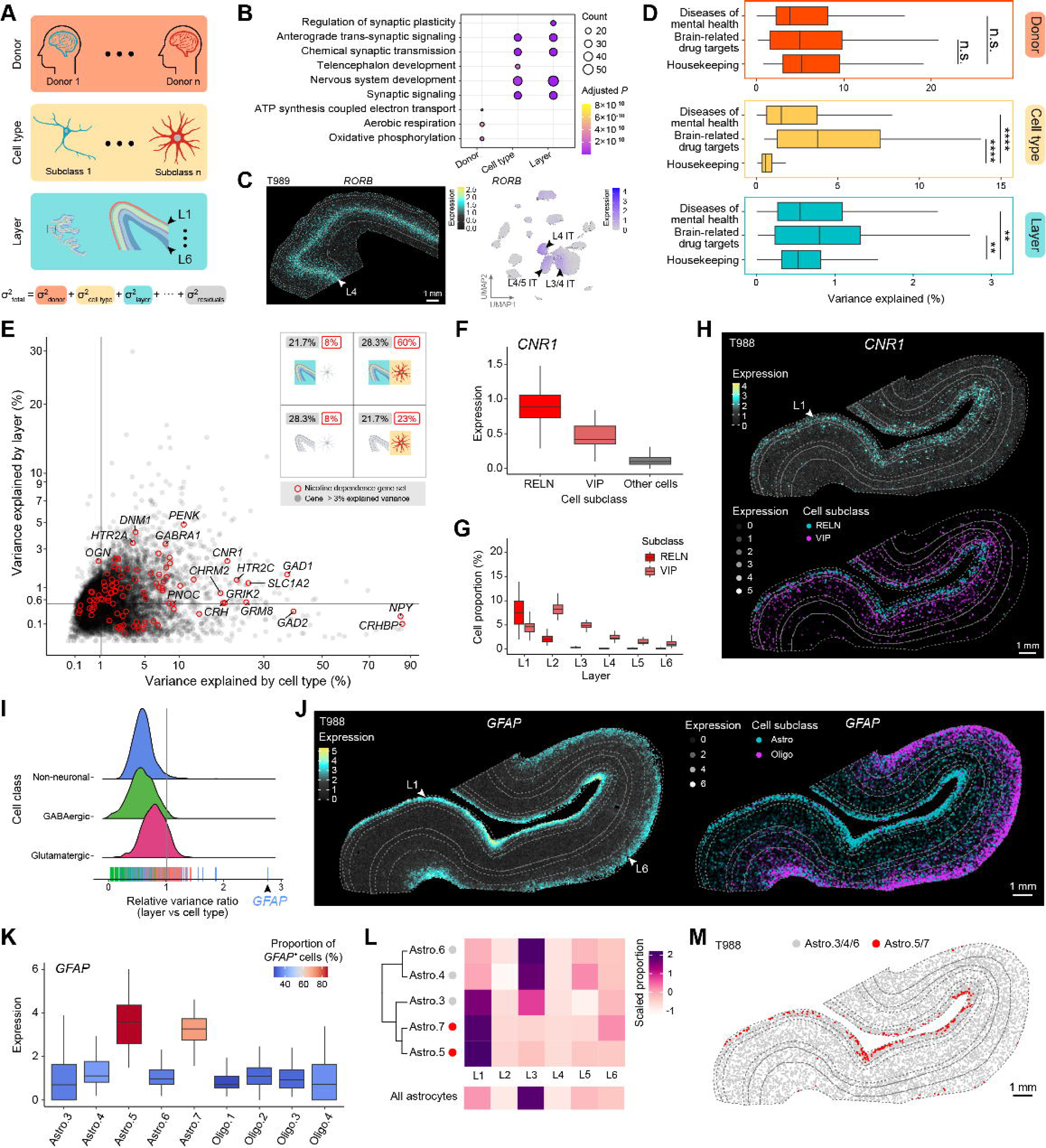
Variance-based multi-factor analysis of gene expression patterns. (**A**) Overview of variance partitioning analysis, highlighting four major sources of gene expression variability: donor, cell type (cell subclass), and cortical layer. (**B**) Dot plot displaying the Gene Ontology (GO) enrichment results of genes with the highest variance (top 100) explained by specific factors. (**C**) Expression pattern of *RORB* gene in a representative Stereo-seq section (right) and snRNA-seq data (left). Scale bar, 1 mm. (**D**) Comparison of explainable variance across disease of mental health genes, brain-related drug target genes, and housekeeping genes for three major factors (donor, cell type and cortical layer). Statistical tests were carried out using Wilcoxon rank-sum test with Benjamini–Hochberg adjustment, with significant levels labeled (* for *P*-value < 0.05, ** for *P*-value < 0.01, *** for *P*-value < 0.001, **** for *P*-value < 0.0001, n.s. for *P*-value ≥ 0.05). (**E**) Scatter plot showing the proportion of variance explained by cell type (x-axis) and cortical layer (y-axis) for all genes. Genes from the nicotine dependence gene set are marked as red circles. All other genes are shown in light gray. The quadrant schematic in the top-right corner classifies genes into four quadrants (Q1, Q2, Q3 and Q4) based on the median of variance. Q1 includes genes with high variance in both cell type and layer; Q2 and Q4 represent genes with high variance in either layer or cell type, respectively; and Q3 contains genes with low variance in both factors. The colored ratios indicate the proportions of genes from the nicotine dependence gene set (red) and all genes (black) within each quadrant, respectively. (**F**) Box plot showing the expression levels of *CNR1* across cell subclasses based on Stereo-seq sections, with each data point representing an individual section for statistical analysis. (**G**) Box plot showing the proportions of RELN and VIP neurons across cortical layers based on Stereo-seq sections, with each data point representing an individual section for statistical analysis. (**H**) Spatial expression pattern of *CNR1* in a representative Stereo-seq section (T988), with the overall expression at bin100 (top; 100×100 DNA nanoballs, DNBs) and the expression in RELN and VIP neurons at single-cell level (bottom). Scale bar, 1 mm. (**I**) Density plot of the relative variance ratio (layer vs cell type) for marker genes of cell subclasses, colored by cell classes. Each tick mark (rug) represents a subclass marker gene. The vertical line indicates equal contributions from layer and cell type. (**J**) Spatial expression pattern of *GFAP* in a representative Stereo-seq section (T988), with the overall expression at bin100 (left) and the expression in astrocytes and oligodendrocytes at single-cell level (right). Scale bar, 1 mm. (**K**) Bar plot showing the expression levels of *GFAP* across astrocyte and oligodendrocyte subtypes based on Stereo-seq sections, with each data point representing an individual section for analysis. (**L**) Heatmap showing the scaled proportions of astrocyte subtypes across cortical layers in Stereo-seq sections. (**M**) Spatial distribution of astrocyte subtypes in a representative Stereo-seq section (T988). Astro.5 and Astro.7 are highlighted in red, while Astro.3, Astro.4, and Astro.6 are shown in gray. Scale bar, 1 mm. See also Figure S4.

Among these factors, donor identity accounted for the largest percentage of the variance, followed by cell type, donor age, and layer location. Collectively, these top four factors explained 98.1% of the total variance from all explainable contributing factors (**Figure S4B)**. Notably, this study provides the first quantitative estimate of layer location’s contribution to transcriptomic variation in spatially resolved cortical data, underscoring its previously underappreciated role in shaping gene expression patterns. Genes ranked by the percentage of various contributing factors displayed diverse biological functions. For instance, genes associated with adenosine triphosphate (ATP) and oxidative respiration tended to exhibit higher individual variation, whereas genes involved in synaptic signaling and transmission showed greater cell type or layer-related variation (**Figure 4B**), consistent with previous human brain single cell population scale studies ^44^. Moreover, we found expression of several genes with high percentages of two or more factors contributing to their variance (**Figure S4C**). As expected, marker genes of glutamatergic neurons tended to have higher percentages of the variance attributable to both the cell type and cortical layer location, as compared with those of GABAergic neurons and non-neuronal cells (**Figures S4D** and **S4E**). This is consistent with more distinct laminar distribution patterns found for glutamatergic neurons (**Figure S1H**). For example, *RORB*, a well-known marker gene for IT neurons ^45^ in cortical L4, exhibited substantial variation due to both the cell type and layer location (**Figure 4C**).

Understanding how genetic, cellular, and spatial factors contribute to gene expression variation is critical for uncovering disease mechanisms and developing targeted therapies. To further investigate the biological relevance on various factors contributing to the expression variance, we evaluated the factors’ association with brain diseases and drug-target gene sets. Specifically, we utilized gene sets associated with central nervous system diseases (e.g. cerebral palsy) and diseases of mental health (e.g. bipolar disorder) from DISEASES database ^46^ and brain-related drug targets from Therapeutic Target Database (TTD) ^47^. Overall, gene sets associated with brain diseases exhibited higher percentages of explainable variance as compared with other diseases (brain-irrelevant diseases) for three major factors: donor identity, cell type, and layer location (**Figure S4F**). Furthermore, the gene set associated with mental health and brain-related drug targets showed significantly higher percentages of variance attributable to cell type and layer location than those for the housekeeping gene set ^48^, but not to the donor identity (**Figure 4D**). These findings suggest that while genetic background (donor identity) contributes substantially to expression variance, cell type specificity and spatial organization (layer location) play particularly critical roles in mental health disorders and the mechanisms of brain-targeted therapies.

For instance, the majority of genes within the nicotine dependence gene set, one of mental health-related disease gene sets (**Figure S4G**), exhibited high percentage of variance attributable to both cell type and cortical layer location (**Figure 4E**). This gene set also showed high expression in GABAergic neurons and was enriched in cortical L2/3/4 (**Figure S4H**), aligning with the role of nicotinic acetylcholine receptors (nAChRs) in GABAergic neurons. These receptors mediate dose-dependent nicotine intake ^49–51^ and exhibit layer-specific modulation ^52,53^. Notably, within this gene set, the cannabinoid receptor 1 *CNR1*, a known therapeutic target for various types of addiction ^54^, exhibited high expression in two distinct neuron populations: RELN and VIP neurons (**Figure 4F**). These cell types exhibit distinct laminar distributions: RELN neurons predominantly localize to L1, while VIP neurons are enriched in L2 (**Figure 4G**), contributing to *CNR1*’s layer-specific expression pattern (**Figure 4H**). The cell-type- and layer-specific expression of *CNR1* underscores the power of spatial transcriptomics in uncovering complex regulatory mechanisms in addiction, revealing how discrete neuronal populations, positioned within precise cortical layers, may be differentially involved in reward-related circuits. These findings contribute to our understanding of addiction neurobiology and open new avenues for developing targeted therapies that account for spatial and cellular heterogeneity.

Besides genes with high variance in expression attributable to both cell type and cortical layer location, we noticed that several genes, mostly marker genes of non-neuronal cells, demonstrated high layer-related variance but relatively low cell type-related variance (**Figure 4I**). A notable example is *GFAP* that encodes glial fibrillary acidic protein, which is known to be enriched mainly in astrocytes and immature oligodendrocytes ^55,56^. Among all genes examined for their expression variance, *GFAP* exhibited the highest relative percentage ratio for variance-contributing factor ratio of layer vs cell type (**Figure 4I**; **STAR Methods**), indicating dominant effect of layer location in determining expression variability. Our spatial analysis revealed that *GFAP* was strongly enriched in cortical L1, and expressed at moderate levels in oligodendrocytes in L6, but not in neurons (**Figures 4J**, **S4I** and **S4J**). We also identified two astrocyte subtypes (Astro.5 and Astro.7) with markedly elevated *GFAP* expression, as well as three other astrocyte subtypes (Astro.3, Astro.4 and Astro.6) exhibited relatively low *GFAP* expression levels (**Figure 4K**). Notably, these two high *GFAP* expressing astrocytes were specifically enriched in L1 (**Figures 4L** and **4M**), indicating the presence of specialized astrocyte subtypes in superficial cortical layers. Further differential expression analysis revealed that these L1-astrocyte subtypes were enriched for genes involved in extracellular matrix organization (**Figures S4K** and **S4L**), suggesting their potential role in maintaining the brain–blood barrier ^57^. Together, these findings demonstrate how spatial transcriptomics can dissect both layer-specific and cell-type-specific expression patterns, revealing specialized glial subpopulations with potentially distinct roles in cortical organization and function.

### Spatial specificity of cis-eQTLs in the human cortex

Single-cell spatial transcriptomics provides a powerful approach to elucidate how genetic variants influence gene expression across diverse brain cell types within their native spatial contexts, thereby refining models of brain-related regulatory mechanisms. However, this approach requires population-scale cohorts profiled with single-cell spatial transcriptomics to robustly link genetic variants to expression and spatial context, to construct accurate single-cell regulatory and cell–cell interaction models. To explore the genetic basis of spatial gene expression regulation, we performed cis-eQTL analysis by integrating WGS with Stereo-seq datasets. Gene expression matrices derived from Stereo-seq sections were aggregated at either cell subclass or cortical layer levels, enabling the identification of cell-type and layer specific cis-eQTLs (**Figure 5A**; **STAR Methods**). We identified 725,869 high-quality single nucleotide polymorphisms (SNPs) from WGS data (**STAR Methods**). Multidimensional scaling (MDS) confirmed that our cohort genetically aligns with the East Asian population from the 1000 Genomes Project ^58–60^, ensuring population comparability and genetic background consistency (**Figure 5B**; **STAR Methods**). In total, we identified 3,113 significant cis-QTLs across all cell subclasses and 3,545 across all cortical layers (FDR < 0.05). The number of detected cis-eQTLs was positively correlated with cell abundance, consistent with the previous single-cell eQTL study ^44^. **(Figures S5A**–**S5D**). Among these high confidence cis-eQTLs, we observed a pronounced enrichment of SNPs located near the transcription start site ^18,27^, aligning with known regulatory principles (**Figures 5C** and **S5E**).

**Figure 5.**
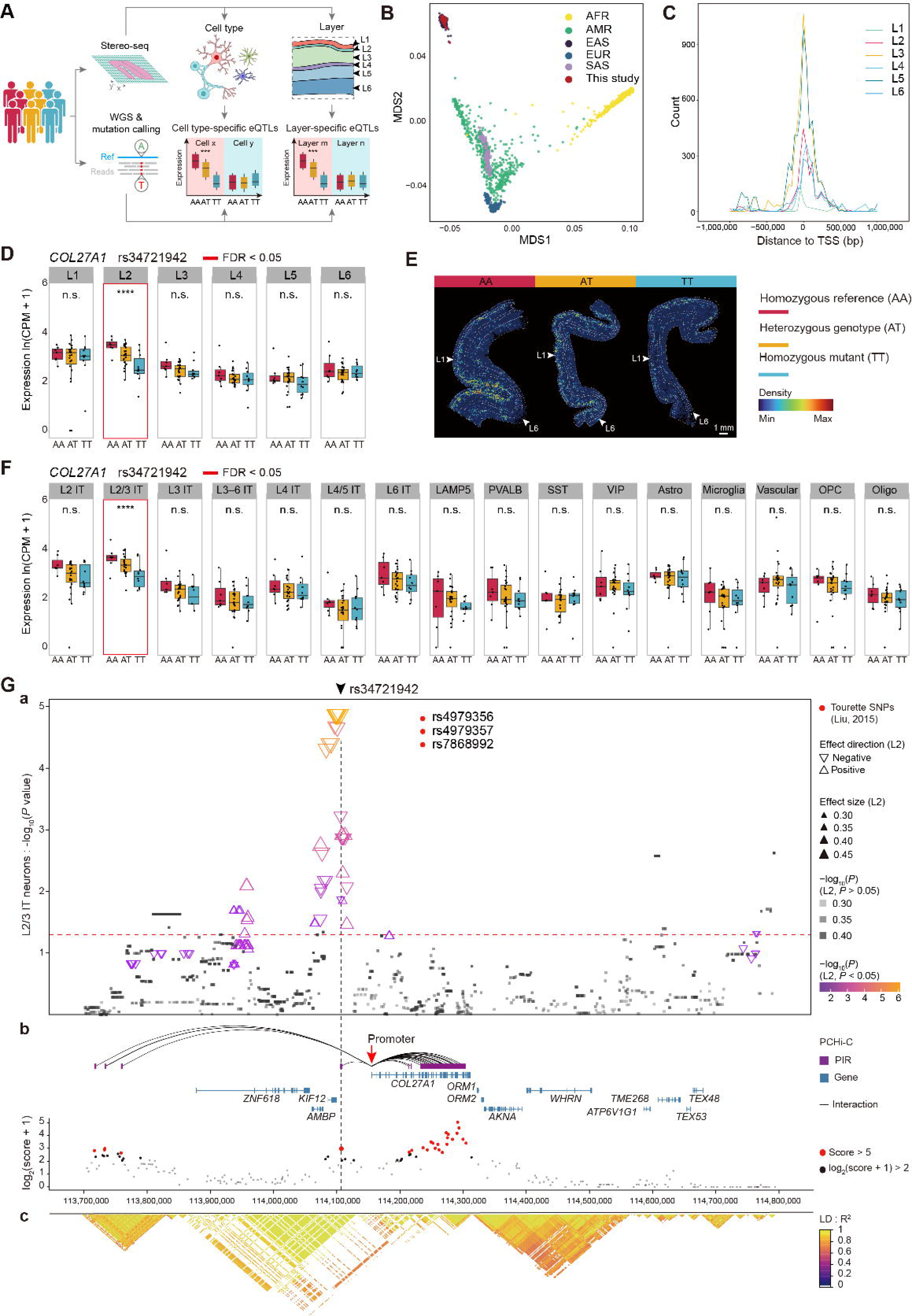
Cell type and layer-specific cis-eQTLs in the human cortex. (**A**) Schematic procedures showing the overview of cis-eQTLs identification workflow. (**B**) Multidimensional scaling (MDS) plot showing the genotyping of our data (colored in red) and samples from the 1000 Genomes Project (AFR, African; AMR, Admixed American; EAS, East Asian; EUR, European; SAS, South Asian). (**C**) Distribution of distances to transcription start sites (TSSs) from significant (FDR < 0.05) cis-eQTLs across cortical layers. (**D**) Box plot showing the expression of *COL27A1* on various rs34721942 genotypes across cortical layers. The significant (FDR < 0.05) layer-specific cis-eQTL is labeled by a red box, with *P*-value labeled (* for *P*-value < 0.05, ** for *P*-value < 0.01, *** for *P*-value < 0.001, **** for *P*-value < 0.0001, n.s. for *P*-value ≥ 0.05). (**E**) The expression density of *COL27A1* on various rs34721942 genotypes in representative Stereo-seq sections (T993, T994 and T736). Highlighting bin100 where *COL27A1* expression is greater than 0, with darker colors indicating a higher local density of such expression-positive bins. (**F**) Box plot showing the expression of *COL27A1* on various rs34721942 genotypes across cell subclasses. The significant (FDR < 0.05) cell type-specific cis-eQTL is marked in red, with *P*-value labeled. (**G**) Integrated views of cis-eQTLs, promoter capture Hi-C (PCHi-C) interactions, and linkage disequilibrium (LD) surrounding *COL27A1*. (**a**) Distribution of SNPs surrounding *COL27A1* (chr9: 113700000-114800000), with a dashed line indicating the position of rs34721942 in the genome. Red dots indicate the position of intronic *COL27A1* SNPs associated with Tourette syndrome. The y-axis displays the significance level of cis-eQTLs related to *COL27A1* in L2/3 IT neurons, with a red horizontal line representing the significance threshold (*P*-value = 0.05). Each triangle represents a SNP, colored by the significance levels of cis-eQTLs related to *COL27A1* in cortical L2, with size representing effect size and direction indicating the effect of regulation. (**b**) Distribution of chromatin interactions from PCHi-C data. The position of *COL27A1* and its neighboring genes on the genome are shown in the blue track. High-confidence PIRs (CHiCAGO score > 5, red dots) are shown in the purple track, connected with *COL27A1* promoter. (**c**) Heatmap showing the pairwise LD measurements between SNPs, with the heatmap colors indicating the levels of LD. See also Figure S5.

To further investigate the layer location and cell-type specificity of genetic regulation, we identified 1,143 cis-eQTL (affecting 172 eGenes) were solely detected in a single cell subclass, and thus defined as cell-type specific cis-eQTLs. For example, the expression of *FAS* significantly associated with the rs11202918 variant specifically in astrocytes (**Figure S5F**). Meanwhile, 1,276 cis-eQTL (affecting 164 eGenes) which were exclusively detected in a single cortical layer, were defined as layer specific cis-eQTLs. As expected, layer-specific eGenes were significantly overlapped with cell-type specific eGenes (**Figure S5G**), consistent with the laminar distribution patterns of cell types in the human cortex. A representative example is *CHRM5*, which encodes the M5 muscarinic acetylcholine receptor and has been proposed as a risk gene involved in schizophrenia and addiction-related pathways ^61,62^. *CHRM5* expression was significantly associated with the SNP rs668922 specifically in cortical L6 (**Figure S5H**). Notably, this SNP also showed a specific association with *CHRM5* expression in oligodendrocytes (**Figure S5I**), a cell type enriched in L6, thereby contributing to the observed layer specificity. Our data showed that individuals homozygous for the mutant allele of rs668922 exhibited markedly reduced *CHRM5* expression in L6 (**Figures S5J** and **S5K**). These findings are consistent with a previous report of an oligodendrocyte-specific eQTL regulating *CHRM5* ^18^, further supporting that its expression is subject to both layer and cell type-specific regulation.

Another cis-eQTL with both laminar and cell-type specificity is rs34721942, a variant located upstream of *COL27A1*, which shows higher allele frequency in East Asian populations according to the dbSNP database ^63^. This SNP significantly affected *COL27A1* expression in cortical L2 and in L2/3 IT neurons, with no detectable effect in other layers or cell subclasses (**Figures 5D**–**5F**), suggesting a layer and cell type-specific genetic regulation. Previous studies have implicated variants related to the *COL27A1* gene in the pathogenesis of various brain-related disorders. For example, SNPs such as rs1588550383 and rs5900078 have been linked to steel syndrome ^64^, a congenital skeletal disorder. In addition, genome-wide association studies (GWAS) of Tourette syndrome, a neurodevelopmental disorder which often co-occurs with psychiatric disorders such as schizophrenia, have identified significant associations with multiple intronic variants in *COL27A1*, such as rs4979356, rs4979357 and rs7868992, indicating the potential role of these variants in the genetic architecture of neuropsychiatric conditions ^65,66^. To further investigate the regulatory mechanisms of these variants, we integrated our cis-eQTL data with promoter capture Hi-C (PCHi-C) data ^67^, enabling the mapping of chromatin interactions surrounding the *COL27A1* gene promoter region (**Figure 5G**). This analysis revealed that significant cis-eQTLs in both cortical L2 and L2/3 IT neurons, such as rs34721942, were predominantly enriched within the promoter-interacting region (PIR), suggesting that these SNPs are in close three-dimensional proximity to the *COL27A1* promoter, potentially facilitating long-range gene expression regulation through direct chromatin interactions ^68^.

Moreover, we observed a dynamic cis-eQTL, rs235331, that exhibited a gradient in effect size along the cortical depth. This SNP predominantly regulated the expression of the immune-modulating gene *AIRE* in superficial layers ^69^, with the effect size progressively decreasing from L2/3 IT to L4/5 IT and L6 IT neurons (**Figures S5L** and **S5M**), showing that the immune-related gene in superficial layers may be more sensitive to genetic regulation. Our findings align with a previous study ^44^ that also reported dynamic eQTLs varying along cortical depth, further supporting the existence of gradient genetic regulation mechanisms in the human cortex.

### Comparative analysis of brain cortices across human, macaque and mouse

The evolutionary expansion of the neocortex is a pivotal event underlying the emergence of higher cognitive functions in primates, with notable differences between the human cortex and that of other mammals highlighted by previous studies ^17,70^. To further investigate species-specific differences in spatial molecular architecture, we performed variance partitioning analysis on Stereo-seq data from the four human cortical regions, including superior frontal gyrus (SFG), middle temporal gyrus (MTG), superior parietal cortex (SPC), and primary visual cortex (V1), integrating homologous regions from macaque and mouse cortex based on our previously generated Stereo-seq dataset ^20,21^ (**Figure 6A**). Our analysis revealed that interspecies differences accounted for a greater proportion of gene expression variance than differences attributable to cortical layers or regions, which demonstrates that evolutionary divergence exerts a more dominant influence on the spatial molecular architecture of the neocortex. Genes with high variance related to species (909 genes, species-associated variance > 20%) were further divided into groups based on the expression levels in various species (**Figure S6A**; **STAR Methods**). GO enrichment analysis indicated that genes highly expressed in humans, or in both humans and macaques, were predominantly associated with synapse organization and postsynaptic specialization (**Figure S6B**). For example, *NPTX1*, one of neuronal pentraxin gene involved in homeostatic synaptic plasticity ^71,72^, showed higher expression in the human cortex compared to the cortical regions of macaque and mouse (**Figure S6C**). Further quantification of differentially expressed genes across species revealed transcriptomic differences in cortical layers and regions.

**Figure 6.**
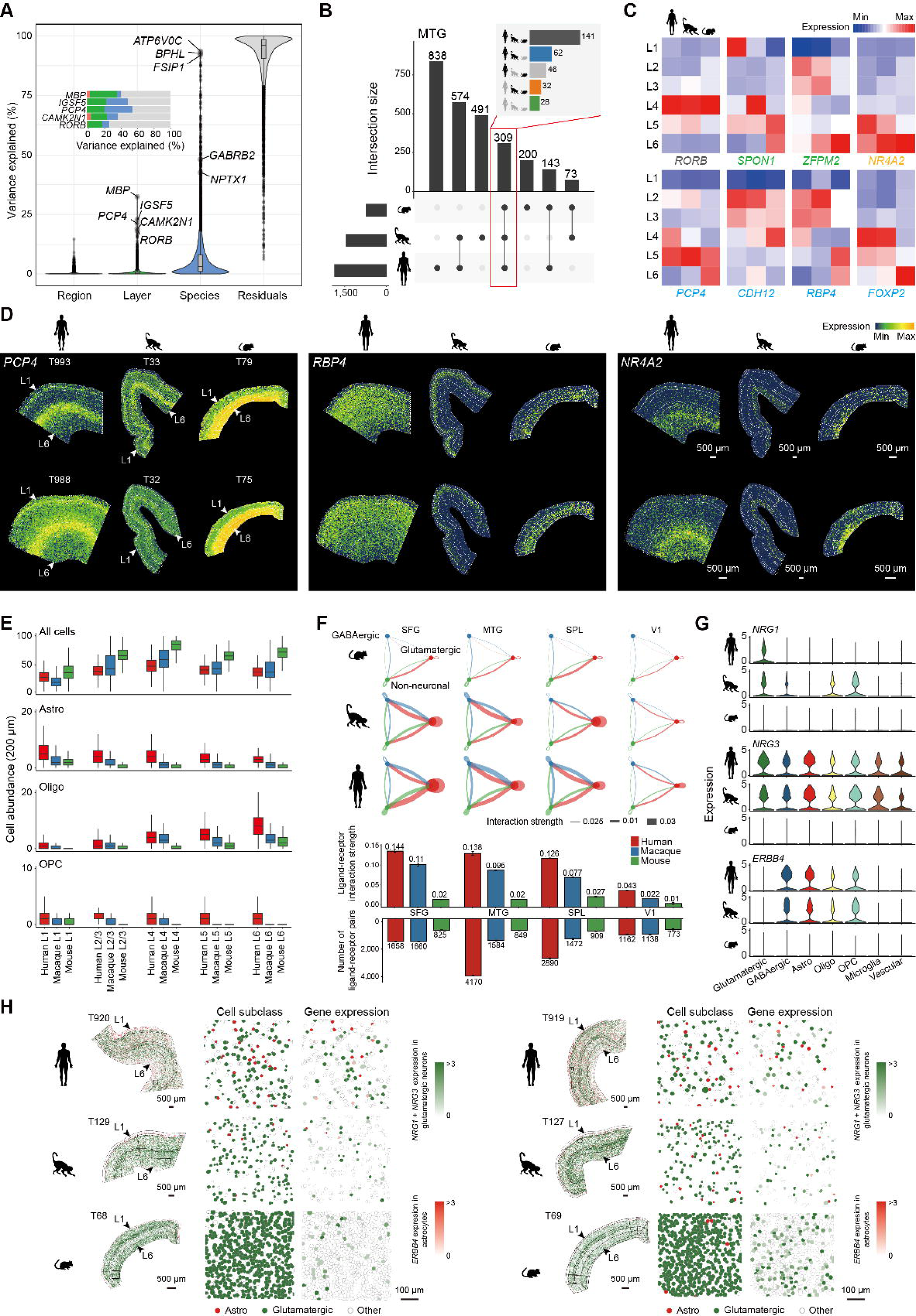
Inter-species differences in gene expression, cellular composition, and intercellular communication in the cortex. (**A**) Violin plot showing the proportion of variance in gene expression explained by annotated factors (cortical regions, layers and species). Representative genes of each factor are highlighted. The top five genes with the highest layer-related variance are emphasized in the bar plot. (**B**) UpSet plot illustrating the intersection of layer-enriched genes in the MTG regions across three species. The x-axis denotes intersection groups, and the y-axis indicates the number of genes per group. The consistency of 309 layer-enriched genes shared across all three species is highlighted in the bar plot, showing the number of genes with conserved or divergent laminar localization patterns across species. (**C**) Distribution of 8 representative genes across layers in different species. Genes are color-coded based on their conservation across species (Black, conserved in 3 species; Green, divergent in 3 species; Orange, conserved in macaque and mouse; Blue, conserved in human and macaque). (**D**) Spatial expression patterns of *PCP4*, *RBP4* and *NR4A2* in the cortex of human, macaque, and mouse, as visualized in representative Stereo-seq sections. Scale bar, 500 µm. (**E**) Box plot showing the abundance of all cell types and specific cell subclasses around glutamatergic neurons across cortical layers. (Astro, astrocyte; Oligo, oligodendrocyte; OPC, oligodendrocyte precursor cell). Cell abundance was quantified by the number of cells within a 200 µm radius. (**F**) The cell–cell interactions between cell classes across species and cortical regions. Chord diagrams illustrating the interaction strength between cell classes. Bar plot showing overall interaction strength and the number of detected ligand–receptor interactions. (**G**) Violin plot illustrating the expression levels of three NRG pathway genes (*NRG1*, *NRG3*, and *ERBB4*) in glutamatergic neurons, GABAergic neurons, and subclasses of non-neuronal cells in the MTG region based on Stereo-seq data. (**H**) Spatial distribution of glutamatergic neurons and astrocytes with highlighted expression of NRG pathway genes. Glutamatergic neurons are shown in red, astrocytes in green, and other cell types are uncolored. The combined expression of *NRG1* and *NRG3* in glutamatergic neurons is represented with a color gradient from white to green, while *ERBB4* expression in astrocytes is displayed with a gradient from white to red. Scale bar, 500 µm; zoom-in regions, 100 µm. **See also Figure S6.**

Among the four cortical regions, the SFG, superior part of the prefrontal cortex, demonstrated the greatest interspecies differences, particularly in cortical L2–5. Otherwise, the V1 showed minimal differences across species (**Figure S6D**), consistent with its conserved functional role in visual processing across mammals ^73,74^, indicating that the transcriptional profile of V1 has been evolutionarily preserved to support fundamental sensory functions.

Notably, we observed that genes with high layer-related variance simultaneously exhibited a high proportion of species-related variance (**Figure 6A**). To further elucidate the conservation and divergence of layer-enriched gene expression across species, we identified layer-enriched genes in corresponding cortical regions of macaque and mouse (**STAR Methods**) and compared them with human layer-enriched genes across cortical lobes. Our analysis revealed a substantial number of genes with conserved laminar specificity across all three species, followed by a notable proportion of genes exhibiting conservation between the two primates, consistently observed in each of the four cortical regions (**Figures 6B** and **S6E**). For instance, 309 genes displayed layer-enriched expression in MTG region across all three species, among which 141 genes exhibited conserved enrichment in the same cortical layer across three species, such as *RORB*, a canonical marker of cortical L4, consistently enriched in L4 across species (**Figures 6C** and **S6F**). In contrast, 28 genes demonstrated totally divergent laminar distribution across three species, such as *SPON1*, enriched in distinct cortical layers across species (**Figures 6C** and **S6F**). Notably, 62 genes showed consistent layer-enriched expression in MTG region between human and macaque but not in mouse, such as *PCP4* and *RBP4*, which were enriched in deep layers in mouse but shifted to relative superficial layers in primates. Moreover, 32 genes showed conservation between macaque and mouse but differed in human. For example, *NR4A2*, a gene coding a nuclear receptor involved in neuronal differentiation ^75,76^, was enriched in L5 in human but in L6 in macaque and mouse (**Figure 6D**). These findings reveal both conserved and species-specific patterns of laminar gene regulation, indicating the neocortical expansion during evolution may be accompanied by the reorganization of laminar gene expression. Together, these results illustrate a complex interplay of deeply conserved and recently divergent laminar gene programs, suggesting that neocortical expansion during evolution was facilitated by the reorganization of transcriptional spatial architecture.

To assess species-specific differences in cellular composition, we quantified the proportions of cell types across four cortical regions in human, macaque and mouse, based on Stereo-seq datasets. Our analysis demonstrated that glutamatergic neurons were present at relatively lower proportions in human compared to macaque and mouse, particularly corticothalamic neurons in cortical L6. In contrast, non-neuronal cells, including astrocytes, oligodendrocytes and oligodendrocyte precursor cells (OPCs), were present in higher proportions in human (**Figure S6G**). These cellular composition differences were consistent across the four cortical regions. Further spatial quantification revealed that, although mouse glutamatergic neurons were surrounded by more cells due to higher overall cell density, human glutamatergic neurons were more frequently surrounded by astrocytes, oligodendrocytes, and OPCs (**Figures 6E**, **S6H** and **S6I**). This pattern was consistently observed across cortical layers and regions, suggesting an overall enhancement of glial support in the human cortex.

Beyond cellular composition, we next investigated species-specific differences in intercellular interactions. Quantitative assessment demonstrated that overall cell communication intensity is stronger and more abundant in human cortical regions compared to macaque and mouse (**Figure 6F**; **STAR Methods**). Among the four cortical regions, SFG showed the greatest interaction strength, while MTG showed the highest number of detected ligand–receptor pairs. In contrast, V1 showed the minimal overall cell communication intensity across the four cortical regions, which also demonstrated lower differences across species (**Figure 6F**). Notably, several signaling pathways related to neural development and synaptic plasticity ^77–80^ exhibited greater communication intensity in primates, such as neuregulin (NRG), neurexin (NRXN), and neural cell adhesion molecule (NCAM) signaling pathways (**Figure S6J**). Among them, two neuregulin coding genes, *NRG1* and *NRG3*, were highly expressed in the primate cortex, with *NRG1* showing enrichment in human glutamatergic neurons. Meanwhile, *ERBB4*, the gene coding a receptor tyrosine kinase, was highly expressed in primate GABAergic neurons and non-neuronal cells, particularly in astrocytes, oligodendrocytes, and OPCs (**Figures 6G**, **S6K** and **S6L**). Spatial visualization demonstrated that human glutamatergic neurons were significantly surrounded by more astrocytes, with neuregulin coding genes highly expressed in glutamatergic neurons and *ERBB4* abundantly expressed in astrocytes (**Figures 6H** and **S6M**), indicating a potential mechanism for enhanced neuron–glia communication in the human cortex. This specialized neuron–glia signaling, particularly through the NRG–ERBB4 pathway, could be a key evolutionary innovation in the primate cortex that contributes to the complex functionality of the human brain.

## Discussion

In this study, we present a comprehensive single-cell spatial transcriptomic atlas of the human cerebral cortex, comprising over 3.4 million spatially resolved cells, covering 34,949 protein-coding and long noncoding RNA genes. By integrating Stereo-seq with snRNA-seq and WGS across diverse donors, brain regions, and age groups, we generated a high-resolution and population-scale resource that reveals the intricate interplay among cellular composition, gene expression, and spatial organization. This atlas surpasses previous single-region or lower-resolution efforts, offering a foundational framework for mapping the molecular architecture of the human cortex.

Previous spatial transcriptomic studies on layer-enriched gene (GLEE) identification have primarily focused on one or two cortical regions and were limited by small donor cohorts ^14,29^. Leveraging our population-scale Stereo-seq dataset with full transcriptome coverage, we systematically identified a broader set of GLEEs across the four major cortical lobes. The consistency and diversity of these genes across lobes indicate the conserved and region-specific genetic programs. Lobe-conserved GLEEs, such as *RELN* and *RORB*, the canonical layer markers for L1 and L4 respectively, are key regulators of neuronal migration and specification during early cortical development ^45,81–83^. GO analysis showed their enrichment in core developmental processes, including neuron projection development and glial cell differentiation, supporting their putative role in cortical circuit formation. In contrast, lobe-specific GLEEs were associated with ion channel activity, indicating the regional differences in electrophysiological activity shaped by spatially distinct gene expression ^84^. Additionally, we identified two occipital lobe-specific GLEEs, *EGR2* and *EGR4*, which encode early-response transcription factors, aligning with prior studies of light-induced transcriptional responses in the mouse visual cortex ^31,32^. We further identified that these two genes were enriched in a specific L2 IT subtype, suggesting a potential role for sensory-driven transcriptional programs in the specialization of L2 IT neurons in the visual cortex. These findings warrant further investigation into their functional roles in cortical processing.

Leveraging our large population-scale atlas, we demonstrate through spatially resolved aging analysis how cortical transcriptomes are dynamically remodeled across the human lifespan. We identified 1,814 age-related genes and 9 gene modules (M1–M9), which exhibited both laminar and cell-type specificity, and revealed a decline in neuronal gene programs alongside increasing activation of glial-associated pathways. This complementary relationship may reflect a regulatory mechanism in the aging process, which is consistent with reports that neural injury triggers changes in glial cell states and proliferation, especially astrocytes ^85,86^. Notably, several neuron subtypes, such as L3 IT.6 excitatory neurons and SST.17 inhibitory interneurons, showed marked reductions in both abundance and module gene expression with age. Moreover, one of the down-regulated gene modules (M6), which was enriched in *SST*^+^ interneurons and related to ion transport, was significantly enriched for genetic risk loci of Alzheimer’s disease and major depression. This finding aligns with previous reports of reduced *SST* expression in normal aging and Alzheimer’s disease ^39–41,87,88^. A recent single-nucleus transcriptomic study further confirmed the age-associated reduction of *SST*^+^ interneurons in the human cortex ^89^. Leveraging spatial transcriptomics, our spatially resolved cell–cell communication analysis revealed a progressive decline in somatostatin signaling and its ligand–receptor interactions with neighboring excitatory neurons during aging, suggesting a potential mechanism underlying the loss of local circuit integrity in the aging brain. These spatial signatures of cellular vulnerability may provide new insights for understanding the cellular basis of cognitive aging and neurodegeneration.

Gene expression is influenced by multiple factors, and variance decomposition of single-nucleus transcriptomes has demonstrated that genes with high cell type-associated variability are often linked to neurological disorders and therapeutic targets ^27,43,44^. Given the strong overlap between cell type and laminar location in the cortex, spatially unresolved single-nucleus data cannot clearly distinguish between these two factors, potentially leading to confounded interpretations. Our spatial transcriptome variance partition analysis further clarified that both cell type and laminar position explained a relatively high proportion of the expression variance in genes associated with mental health. For example, *CNR1*, a cannabinoid receptor that serves as a therapeutic target for addiction, was enriched in RELN and VIP neurons in L1 and L2 respectively, indicating the different neuronal circuits in different cortical layers. Moreover, leveraging WGS data and spatially resolved cis-eQTL analysis, we identified thousands of regulatory variants that act in a cell type- or layer-specific manner, revealing signals that would be masked in conventional bulk tissue analysis lacking spatial and cell type information. For example, we identified cis-eQTLs at rs668922 and rs34721942 regulating *CHRM5* in oligodendrocytes (L6) and *COL27A1* in L2/3 IT neurons (L2), respectively. These genes have been previously implicated in schizophrenia and addictions (*CHRM5*), and Tourette syndrome (*COL27A1*). Such context-specific eQTLs may reflect the localized availability of regulatory components within particular cellular and laminar environments. Our findings emphasize the importance of spatial cell-type resolution in understanding disease mechanisms, suggesting that effective therapies, such as pharmacological agents and deep brain stimulation, should target the precise cell types and layer locations where pathological processes occur, rather than uniformly across the cortex.

Compared with other mammals, humans exhibit a rapid and nonuniform expansion of the cerebral cortex, particularly in prefrontal and association regions, reflecting functional specializations integral to higher cognition and social behavior ^74,90–92^. Our cross-species comparisons revealed substantial transcriptomic divergence throughout evolution, with markedly higher interspecies differences in prefrontal and association regions (SFG, MTG, SPC) than in the evolutionarily conserved primary visual cortex (V1). These transcriptomic changes were accompanied by reduced neuronal density, increased glial cell abundance, and enhanced neuron–glia interactions, consistent with classic histological observations that glia-to-neuron ratios rise with increasing brain size and complexity, alongside declining neuronal density ^93–95^. Notably, interaction pathways related to synaptic plasticity and neural development, such as the neuregulin (NRG) signaling pathway ^77–80^, were more active in the human cortex compared with macaque or mouse. Previous studies have shown that NRG signaling enhances neuroprotection via astrocytic glutamine synthetase upregulation ^77^ and oligodendrocyte-mediated remyelination ^78^. Our findings thus underscore the integral role of glia and neuron–glia communication in supporting human-specific cortical functions and may provide mechanistic insights into neuropsychiatric disorders.

In summary, our work provides a spatially resolved single-cell atlas of the human cortex across cortical regions and a broad age range spanning the human lifespan, integrated with individual genetic information, revealing spatial patterns of cell types and gene expression associated with cortical region specialization, aging, regulatory variation, and evolutionary divergence. This atlas not only advances fundamental neuroscience but also offers a valuable resource for decoding disease mechanisms and designing spatially informed therapeutic strategies.

### Limitations of the study

There are several limitations to this study. First, due to the limited capture efficiency of gene transcripts inherent to barcoding-based spatial transcriptomics methods ^23^, the average number of genes captured per cell (533) was lower than that typically achieved with conventional snRNA-seq, and was insufficient for precise cell annotation. Therefore, we used our snRNA-seq data to assist cell-type identification. Second, although thousands of cis-eQTLs were identified in our spatial transcriptomic and WGS datasets, the number remains relatively limited. This likely reflects the modest statistical power at current sample size, and suggests that additional donors would enable the discovery of more spatially resolved regulatory variants. Third, our dataset comprises only East Asian individuals. Further studies are required to assess the consistency of spatial and regulatory patterns in individuals of diverse genetic ancestries.

## STAR methods

### Key resources table

See separate file Key_resource_table.docx.

### Resource availability

#### Lead contact

Further information and requests for the resources and reagents may be directed to and will be fulfilled by the lead contact, Wu Wei.

#### Materials availability

All materials used for stereo-seq and snRNA-seq are commercially available.

#### Data and code availability

All processed data and raw data have been deposited to CNGB Nucleotide Sequence Archive under accession codes CNP0007621 for snRNA-seq data and CNP0009546 for Stereo-seq data (https://db.cngb.org). All data were analyzed with standard programs and packages, as detailed in the key resources table. Additional information required to reanalyze the data reported in this paper is available from the lead contact upon request.

### Experimental model and study participant details

#### Ethical considerations

Human brain tissue was collected from patients who were pathologically diagnosed with tumor, epilepsy or abscess. All experimental samples involved in this study were approved by the Ethics Committee of Huashan Hospital (No. KY2022-590), Children’s Hospital of Fudan University (No. (2023)313), and Shanghai Neuromedical Center (No. DJKY2022-003)., and the Institutional Review Board on Ethics Committee of BGI group (No. BGI-IRB 24104). The normative requirements of the relevant sample manipulation were strictly adhered.

### Method details

#### Brain tissue collection for Stereo-seq

Immediately after human brain tissue is removed during neurosurgery, it is transferred to a disposable petri dish filled with pre-cooled OCT. The OCT is replaced three times to displace the liquid on the surface of the sample and to prevent the formation of surface ice crystals due to uneven heating during snap-freezing. Since it is not possible to guarantee a standard coronal, horizontal, or sagittal plane during surgical resection, the largest surface of the sample is usually used as the embedding base after OCT treatment, placed in a homemade metal embedding frame, and placed on dry ice for quick freezing after submerging the entire tissue sample with OCT. After quick freezing, the samples were transferred to a −80 L refrigerator for storage.

#### Tissue cryosection, section flattening and RNA quality control

Prior to frozen sectioning, the tissue blocks were transferred 1–2 hours in advance from the −80 L refrigerator to the slicer (Thermo Fisher Cryostar NX50) for equilibrium rewarming, with the slicer chamber temperature at −20 L and the freezing head temperature generally controlled between −8 L and −10 L. Before the official start of sectioning, the tissue samples were glued to the sample trays using OCT and fixed to the universal head of the slicer for trimming and removing the OCT from the tissue surface, a process that ensures that flat, full-tissue-block-size slices are obtained before the tissues are exposed to minimize the loss of tissue samples.

Once the tissue was exposed, the slices were continued to cut to the intact cortical layers 1–6 to begin harvesting the slices. This step was mainly to ensure that the slices contained the full cortex of the brain mass at the time of patching to avoid subsequent problems of missing data. The slice thickness was 10 μm, and two consecutive slices were collected for patching after determining the patching position, while one retention slice was kept in front of and behind the chip, and two slices were kept for subsequent immunohistochemical staining verification.

Upon obtaining to the sections for chip patching, the tissue sections were first spread out with a soft bristle brush and plastic tweezers, and then carefully placed on the Stereo-seq chip pre-cooled 5 minutes in advance. After that, the chip with slices attached was transferred to a baking machine for 5 minutes to bake the slices to dry out the water in the tissue, and then the subsequent Stereo-seq experimental process was continued.

In order to ensure the quality of the samples, 1–2 additional 10 μm slices will be collected at the end of the sectioning of each brain block for random Total RNA extraction and quality testing, and the RNA RIN value greater than or equal to 7 is considered as qualified samples. After all the slicing work was finished, if the brain block was not finished, it would be sealed with OCT and put into −80 L refrigerator together with the retained slices for freezing.

#### Stereo-seq experiment procedure

The Stereo-seq protocol was implemented following methodological guidelines reported in the macaque cortex Stereo-seq study ^20^. Stereo-seq microarrays (1 cm × 1 cm or 1 cm × 2 cm) with tissue sections affixed to them were baked at 37°C for 5 minutes, then fixed in methanol (Sigma, #34860, pre-cooled at −20°C for 30 minutes; 24-well plates with 2.5 mL methanol per well) and incubated at −20°C for 30 minutes. At the end of the fixation period, the microarrays were air-dried in a fume hood (to remove the methanol), and stained for 5 minutes with nucleic acid dye (ssDNA reagent, Invitrogen, Q10212) and FITC-conjugated Concanavalin A (ConA, Vector Labs, FL-1001) to stain the tissue sections on the microarrays, and washed with 0.1xSSC buffer (Ambion, AM9770; containing 0.05 U/μL RNase inhibitor) at the end of staining. Fluorescence images were taken using a Zeiss Z1 microscope.

Tissue sections were immersed in permeabilization reagent and incubated at 37°C for 15 minutes (permeabilization reagent was 0.1% pepsin, pre-warmed at 37°C for 3 minutes in advance), and then rinsed with 0.1xSSC buffer (containing 0.05 U/μL RNase inhibitor) to remove pepsin. In this step, RNA was released from the tissue and captured by nanospheres on the Stereo-seq chip. Afterwards, Reverse Transcription Mix equilibrated to room temperature was added dropwise to the chip, and the RNA was reverse transcribed at 42°C for 2 h. After reverse transcription, the tissue sections were washed with 0.1x SSC buffer, digested with tissue removal buffer (10 mM Tris-HCl, 25 mM EDTA, 100 mM NaCl, 0.5% SDS) at 55°C for 30 minutes, then washed twice with 0.1x SSC buffer. The cDNA-containing microarray was treated with exonuclease I (NEB, M0293L) at 37°C for 1 h and then washed once with 0.1x SSC buffer for cDNA amplification. the PCR reaction protocol was: incubation at 95°C for 5 minutes, 98°C for 20 s, 58°C for 20 s, and 72°C for 3 minutes for 15 cycles, followed by incubation at 72°C for 5 minutes. PCR products were purified at 55°C for 30 minutes with 0.6 X VAHTSTM DNA Clean Beads, and quantified by Qubit dsDNA HS kit (Invitrogen, Q32854).

#### Stereo-seq library construction and sequencing

PCR products were purified using VAHTS DNA Clean Beads (Vazyme, N411-03; 0.6× bead-to-sample ratio) and quantified via the Qubit™ dsDNA Assay Kit (Thermo, Q32854). A 20 ng aliquot of cDNA underwent tagmentation with in-house Tn5 transposase (55°C, 10 minutes), followed by reaction termination with 0.02% SDS and gentle room-temperature incubation (5 minutes). For library amplification, 25 μL of fragmented DNA was combined with 1× KAPA HiFi HotStart enzyme, 0.3 μM Stereo-seq-Library-F primer (5’-CTGCTGACGTACTGAGAGGCA-3’), and 0.3 μM Stereo-seq-Library-R primer (5’-GAGACGTTCTCGACTCAGCAGA-3’) in a 100 μL reaction system (NF-HLO-adjusted). Thermal cycling parameters included: 95°C for 5 minutes; 13 cycles of 98°C (20 s), 58°C (20 s), 72°C (30 s); final extension at 72°C for 5 minutes. Amplified libraries were double-purified with AMPure XP Beads (0.6× and 0.15× ratios) and processed into DNA nanoballs (DNBs). Sequencing was ultimately performed on an MGI DNBSEQ-TX platform at China National GeneBank (CNGB).

#### Processing of Stereo-seq raw data

Stereo-seq data processing initiated with coordinate identity (CID) sequence alignment to chip design coordinates, permitting 1-base mismatch tolerance during first-read mapping. Reads containing ambiguous bases (N) or ≥ 2 low-quality bases (Phred score < 10) in molecular identifiers (MIDs) were excluded, followed by CID/MID header annotation. Surviving reads underwent genome alignment (GRCh38.p12) via STAR ^119^ with quality filtering (MAPQ > 10), then gene annotation. Unique molecular identifiers (UMIs) sharing identical spatial coordinates (CID) and genomic loci underwent error-aware consolidation (1-mismatch tolerance) to resolve PCR/sequencing artifacts. This spatial resolution framework generated CID-encoded expression matrices through the SAW pipeline (https://github.com/STOmics/SAW), preserving positional transcriptomic information while mitigating technical noise.

#### Image registration and anatomy-based parcellation

Nucleic acid staining images of tissue sections were automatically registered to the corresponding CID-derived mRNA coordinate systems using periodic track lines pre-engraved in the chip planes. For registration, the captured mRNAs of each section were first encoded into a 16-bit image by converting the UMI counts at each coordinate into pixel values. 1D Laplacian of Gaussian (LoG) operators (sigma = 5×10^-4^) were applied independently along the row and column directions of each paired nucleic acid staining image and mRNA image to detect track lines. The relative rotation between two images was estimated by measuring the angular difference between their track lines. After rotation correction, the relative scale and translation were computed using RANSAC algorithm.

When determining the brain region and cortical layers, firstly, the brain region of the sample is identified based on the patient’s brain MRI positioning during sampling. During the manual division of the cortex, immunohistochemical staining was performed on the corresponding frozen sections of the chip. Take the sealed frozen sections out of the −80L refrigerator and allow them to thaw at room temperature (after thawing, open the seal and warm to room temperature, then remove excess OCT; use a histology pen to delineate the reaction area. After that, fix in 4% PFA for 10 minutes, then wash three times with 0.01M PBS (pH 7.4); incubate in 0.5% PBS (Triton X-100 in 0.01M PBS) at room temperature for 15 minutes, then continue incubating at room temperature with 10% serum for 1 hour. Next, mix the primary antibody (NeuN:1/1500) with 0.1% PBST (pre-chilled) and incubate at 4°C overnight; the next day, recover the primary antibody and wash the sections three times with 0.01M PBS; continue incubation with 0.6% H_2_O_2_ (prepared in 0.01M PBS) for 30 minutes, and after that, wash three times with 0.1% PBST, each wash lasting 5 minutes; use the secondary antibody (prepared in 0.1% PBST) to incubate at room temperature for 2 hours, and then continue to wash three times with 0.01M PBS. Equipped with ABCkit (1 drop A and 1 drop B: 5 mLPBS) incubate brain slices at room temperature for 2 h, and rinse once with 0.01 M PBS and 0.05 M tris twice after the end; Add 0.05 B dropwise for 10 minutes, then add 0.05% H_2_O_2_ for color development, about 10 minutes, rinse with 0.05 M Tris (pH 7.6) for 3 times after color development to stop the reaction (avoid light during DAB staining until the reaction is stopped): dehydrate with 50%, 70%, and 100% alcohol gradients, then soak with toluene for 5 minutes, and use DPX gel to dry the gel after two soaking for scanning and imaging. Finally, based on the immunohistochemical images of the corresponding brain slice locations and the differences in cell density across different layers, the cortical layers 1–6 and the white matter area of the brain slice were manually delineated, ultimately resulting in layered results based on anatomy.

#### Screening of high-quality Stereo-seq sections

Stereo-seq sections were defined as high quality if they met the following criteria: (1) In bin100 units (100 DNB × 100 DNB), the average number of detected genes exceeded 1000, and the proportion of mitochondrial gene expression was greater than 25%. (2) The section exhibited clearly delineated cortical layers from L1 to L6 in the BayesSpace ^120^ clustering results. Only the gray mater regions of high-quality sections were used for downstream analysis.

#### Cell segmentation of Stereo-seq sections

The same strategy as described in the macaque cortex Stereo-seq study ^20^ was utilized to segment single cells in registered nucleic acid staining images of Stereo-seq sections. Concanavalin A (ConA) staining images, along with corresponding nucleic acid staining images, were used to generate cell annotations through an active learning strategy. Since ConA stains the cell membranes and provides clear boundaries between cells, it facilitates easier distinction and manual annotation. To begin, a small number of clearly identifiable cells in the ConA staining images were manually annotated and used to train a mask R-CNN-based deep learning model, referred to as the ConA model. The trained ConA model was then applied to unlabeled ConA images to generate preliminary annotations. The generated annotations were manually reviewed, corrected, and next fed back to further fine-tune the ConA model. Sufficient annotations were obtained through the active learning procedure over several iterations, minimizing human involvement, and subsequently transferred to the corresbonding nucleic acid staining images. The nucleic acid staining images, now with transferred cell annotations, were used to train another mask R-CNN-based deep learning model, referred to as the nucleic acid model. Finally, the well-trained nucleic acid model was deployed as the ultimate cell segmentation model to perform single cell segmentation on nucleic acid staining images of all Stereo-seq sections.

#### Sample preparation for single-nucleus sequencing

There are two main methods we use when performing single-nucleus sampling:

1. Whole brain block slicing method: The brain blocks previously used for spatial transcriptome experiments were retrieved from the −80 L refrigerator again and put in the slicer to equilibrate and rewarm. Slices of 70-100 μm thickness were collected from the same anatomical location as used in spatial transcriptomics. Roughly 7-10 slices were collected from each brain block, and single-cell slices from the same brain block were placed in the same freezing tube. Immediately after the collection was completed, the cryostat tubes were tightened and labeled, followed by rapid treatment with liquid nitrogen and storage in a −80 L refrigerator. This method is suitable for larger tissue samples, and helps preserve the cellular diversity of the sampled brain region by encompassing wider cross-sections.
2. Fine cortical sampling method: After removing the OCT from the surface of the brain block, a pre-cooled circular sampler was used to take fine samples of the cortical part of the brain block. Depending on the diameter of the circular hole of the sampler, a block of tissue of 3 mm × 3 mm × 1 mm in size can be obtained at one time. In order to ensure a sufficient number of nuclei for single-tube sequencing, 3 blocks of tissue were taken from each brain block and placed in the same freezing tube, and the subsequent operation is consistent with section sampling. This method is to ensure that the sampling area should encompass layers 1-6 of the cortex, which is generally suitable for smaller volume tissue samples. It not only preserves layer-specific cellular content, but also facilitates the alignment of single-nucleus sequencing data to the corresponding spatial transcriptomic data. In addition, this strategy avoids contamination from white matter.

#### Single-nucleus suspension preparation

Following previously established protocols ^21^, frozen brain tissues were quickly transferred to a 1 mL Dounce homogenizer (TIANDZ) and homogenized in 1 mL of homogenization buffer. The resulting solution was then filtered through a cell strainer into a 1.5 mL tube (Eppendorf), followed by centrifugation at 500 g for 5 minutes at 4 ℃ to pellet the nuclei. The pelleted nuclei were subsequently resuspended in buffer and adjusted to a final concentration of 1000 nuclei/μL for downstream library preparation.

#### Single-nucleus library preparation and sequencing

Single-nucleus libraries were constructed using the DNBelab C Series Single-Cell Library Prep Set (MGI, 1000021082) following established protocols ^21^. The workflow involved generating barcoded libraries through droplet-based encapsulation, emulsion disruption, bead recovery, reverse transcription, and cDNA amplification. Library quantification was performed with a Qubit ssDNA Assay Kit (Thermo Fisher, Q10212) prior to sequencing on a DNBSEQ-Tx platform (China National GeneBank, Shenzhen) under a paired-end configuration (Read 1: 41 bp; Read 2: 100 bp). All experimental steps adhered to manufacturer-recommended specifications.

#### Single-nucleus raw data processing

Processed DNBSEQ-Tx sequencing data were mapped to the GRCh38.p12 reference genome. Given that single-nucleus RNA sequencing (snRNA-seq) inherently captures nuclear pre-mRNA containing unspliced intronic regions, gene expression quantification incorporated both exonic and intronic reads to maximize transcriptomic coverage. Ambient RNA contamination was computationally corrected using SoupX (v1.6.2) ^117^ with tissue-specific background noise estimation via the autoEstCont function for parameter optimization. This dual read-inclusion strategy ensures comprehensive detection of nascent transcriptional activity while mitigating technical artifacts.

#### Quality control of snRNA-seq data

Across all single-nucleus RNA sequencing (snRNA-seq) libraries, those with a median gene count ≤ 700 were excluded, resulting in 114 high-quality libraries retained for downstream analysis. At the nucleus level, doublets were identified and removed using DoubletFinder (v2.0.4) ^118^. Further quality control was applied by filtering out nuclei with the median gene count ≤ 500, mitochondrial gene content ≥ 10%, or UMI-to-gene count ratio ≤ 1.1. Gene expression data were normalized using the NormalizeData function in the Seurat package (v5.0.0) ^98^, with the “LogNormalize” method, followed by dimensionality reduction via principal component analysis (PCA) using the RunPCA function. For library integration, we employed a customized version of the RunHarmony function from the Harmony package (v0.1.0) ^100^. Specifically, we replaced the default k-means clustering algorithm (Hartigan-Wong) with the Lloyd algorithm by introducing a new kmeans.algorithm parameter, along with kmeans.iter.max and kmeans.nstart to accommodate large-scale datasets. The integration was performed with the following settings: assay.use = “RNA”, reduction = “pca”, dims.use = 1:30, kmeans.algorithm = “Lloyd”, kmeans.iter.max = 1000, kmeans.nstart = 10, epsilon.cluster = -Inf, and epsilon.harmony = -Inf. Unsupervised clustering was carried out using Seurat’s FindNeighbors and FindClusters functions (parameters: dims = 1:30, reduction = “harmony”). Based on canonical marker gene expression, clusters were annotated into 7 major cell types: glutamatergic neurons, GABAergic neurons, astrocytes, oligodendrocytes, oligodendrocyte precursor cells (OPCs), microglia, and vascular cells. Nuclei that could not be confidently assigned to any of these types due to ambiguous marker expression were labeled as uncertain nuclei.

During the initial cell type annotation based on clustering and marker gene expression, a subset of nuclei could not be confidently assigned to any of the seven major types due to ambiguous or weak marker gene expression. These nuclei were temporarily labeled as uncertain nuclei. To improve the overall data quality, we trained a random forest classifier to distinguish well-defined nuclei from these uncertain cases. For model training, we downsampled each major cell type, selecting up to 10,000 nuclei (capped at 80% of the total nuclei per type) as the training set. Marker genes for each class were identified using Seurat’s FindAllMarkers function, and the top 100 genes ranked by fold change were selected as features. The random forest classifier was trained using the randomForest function from the randomForest package (v4.7.1.1) ^99^, with log-normalized gene expression values as input and ntree = 500. The trained model was then applied to all nuclei using the predict function, assigning each nucleus to the cell type with the highest predicted probability. Nuclei with a maximum prediction probability ≤ 0.6 were excluded from further analysis. Following iterative clustering and annotation (described below), clusters lacking clear marker gene signatures were also removed. After all filtering steps, a total of 458,914 high-quality nuclei were retained for downstream analyses.

#### Iterative clustering and classification of snRNA-seq data

After filtering and annotating the 7 major cell types, nuclei were further classified into 25 cell subclasses. To achieve finer-resolution clustering, we applied an iterative clustering strategy ^21^ adapted from the scrattch.hicat package (https://github.com/AllenInstitute/scrattch.hicat), in which batch effects across libraries were corrected at each iteration using the Harmony algorithm. Starting from the 25 initial subclasses, we applied the IterCluster recursive function to each subclass individually, which generated a set of progressively refined clusters. To avoid over-splitting, the MergeCluster function was used to merge clusters with insufficient differences in gene expression. Final cluster annotations were based on subclass-specific marker genes, and clusters lacking clear marker gene signatures were excluded. This iterative process resulted in the identification of 248 cell subtypes, comprising a total of 458,914 high-quality nuclei.

#### Independence test of snRNA-seq cell subtypes

To assess the independence of clusters derived from the iterative clustering process, we evaluated whether each cluster exhibited sufficient transcriptional distinction from the others. A random forest classifier was constructed using the randomForest package (v4.7.1.1) to distinguish all 248 cell subtypes. Marker genes for each cluster were identified using the FindAllMarkers function in the Seurat package. Only genes with the adjusted *P*-value (p_val_adj) < 0.05 and average log_2_ fold change (avg_log_2_FC) > 0 were retained. For each cluster, the top 50 genes ranked by avg_log_2_FC were selected as features for model construction. To ensure balanced training, 200 nuclei were randomly downsampled from each cluster. The log-normalized gene expression values were used as input for training the random forest model (ntree = 1000). Model performance was assessed using the out-of-bag (OOB) confusion matrix, and the resulting prediction accuracy was used as a measure of the independence and distinctiveness of each cell subtype.

#### Co-clustering of snRNA-seq data with published datasets

We incorporated two publicly available single-nucleus RNA-seq (snRNA-seq) datasets as references. One dataset represents a whole-brain human snRNA-seq atlas ^8^, while the other includes nuclei derived from specific cortical regions (the middle temporal gyrus and frontal cortex) ^27^. From each dataset, we extracted nuclei collected from the human cerebral cortex and removed any clusters containing fewer than 20 nuclei to ensure reliable comparison.

For each dataset, we randomly downsampled 200 nuclei per cell type cluster. Data normalization was performed using the NormalizeData function from the Seurat package (parameters: normalization.method = “LogNormalize”, scale.factor = 10000). Marker genes for each cell subtype were identified using the FindAllMarkers function, retaining only genes with an adjusted *P*-value (p_val_adj) < 0.05 and an average log_2_ fold change (avg_log_2_FC) > 0. For each cluster, the top 50 marker genes ranked by avg_log_2_FC were selected as the marker gene set. Principal component analysis (PCA) was then performed on each dataset using the RunPCA function (parameters: dims = 1:50), based on the selected marker genes. We applied the RunHarmony function to integrate our dataset with each of the public datasets (parameters: assay.use = “RNA”, reduction = “pca”, dims.use = 1:30, kmeans.algorithm = “Lloyd”, kmeans.iter.max = 1000, kmeans.nstart = 10, epsilon.cluster = -Inf, epsilon.harmony = -Inf). After integration, clustering was performed using the FindNeighbors (parameters: reduction = “harmony”, dims = 1:50) and FindClusters (parameters: resolution = 2) functions.

To evaluate the correspondence between cell types across datasets, we calculated the proportion *R* of a cell type *x* in our dataset that co-clustered with a cell type *y* from a reference dataset using the following formula:

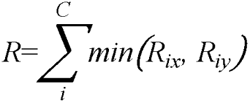

where *C* denotes the set of all clusters obtained from the integrated clustering, *R_ix_* represents the proportion of cell type *x* in cluster *i* relative to the total number of nuclei of type x, and *R_iy_* represents the corresponding proportion for cell type *y*.

#### Stereo-seq cell type transfer

We used Spatial-ID ^28^ to annotate the preserved cells with cell types defined in the snRNA-seq data. The cell type transfer process of Spatial-ID mainly included two stages. In the first stage, deep neural networks (DNN), consisting of four linear layers and the corresponding activation functions, were employed to filter the snRNA-seq data and subsequently trained on the filtered data. The top 100 marker genes for each cell subtype, sorted by fold changes and excluding those associated with mitochondria or ribosomes, were calculated and used as input features for the DNN models. We used a five-fold cross validation strategy to go through the entire snRNA-seq data, retaining only the cells with predicted probabilities larger than 0.5. A DNN was then trained on the filtered snRNA-seq data and subsequently applied to the Stereo-seq data to obtain initial predictions. The second stage involved a graph convolutional network (GCN) consisting of a deep autoencoder, a variational graph autoencoder and a linear classifier. The GCN model incorporated gene expression profiles and spatial neighborhood information, represented by an adjacency matrix, to refine the initial predictions and produce final predicted probabilities. The cell subtype corresponding to the highest predicted probability was ultimately selected as the cell subtype for each cell in the Stereo-seq data.

#### Evaluation of cell type transfer accuracy

To evaluate the accuracy of cell type transfer using Spatial-ID, we computed the scaled Euclidean distance similarity between cell types annotated in Stereo-seq sections and those identified from snRNA-seq data, based on marker gene expression profiles. Marker genes were identified for each cell subclass or cluster using the FindAllMarkers function, with selection thresholds set at *P*-value < 0.05 and average log_2_ fold change (avg_log_2_FC) > 0. The top 500 marker genes, ranked by avg_log_2_FC, were selected as features for similarity calculation. The Euclidean distance similarity score S was defined as:

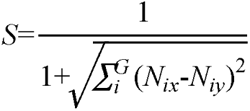

where *N_ix_* and *N_iy_* represent the log-normalized average expression levels of gene i in the Stereo-seq and snRNA-seq datasets, respectively, and G denotes the set of selected marker genes. To ensure comparability across different cell types, the similarity scores were standardized using Z-score normalization. For cell subtypes, similarity scores were scaled within each of the three cell classes.

#### Layer-enriched expressed genes classification

We used the FindAllMarkers function from Seurat package to identify differentially expressed genes across layers (L1–6) in each section. By applying thresholds of adjusted *P*-value (p_val_adj) < 0.05 and average log_2_ fold-change (avg_log_2_FC) > 0.15, we defined genes with cortical layer-enriched expression (GLEEs) for each section. Next, sections were grouped by cortical lobe, and we calculated the frequency of each gene’s appearance as a GLEE across all sections within the same cortical lobe and layer. Genes observed in > 50% of sections within a cortical lobe and layer were defined as GLEEs for that specific cortical lobe and layer. Overall, we identified 906 GLEEs.

GLEEs were classified into three categories (**Figure 2A**): (1) 419 lobe-conserved GLEEs (C-GLEEs), showing consistent laminar expression across all four cortical lobes. These were subdivided into: 255 single-layer C-GLEEs (1L-C-GLEEs), 145 two-layer C-GLEEs (2L-C-GLEEs) and 19 multi-layer C-GLEEs (≥ 3L-C-GLEEs). (2) 406 lobe-specific GLEEs (S-GLEEs), classified by the number of lobes with laminar enrichment: 131 single-lobe-specific S-GLEEs (1Lo-S-GLEEs), subdivided into four subtypes: Frontal lobe (FL-S-GLEEs), Temporal lobe (TL-S-GLEEs), Parietal lobe (PL-S-GLEEs), and Occipital lobe (OL-S-GLEEs). 154 two-lobe-specific S-GLEEs (2Lo-S-GLEEs) and 121 three-lobe-specific S-GLEEs (3Lo-S-GLEEs). (3) 81 Others-GLEEs, exhibiting pan-lobar expression (across all four lobes) but inconsistent layer-enrichment patterns between cortical lobes.

#### Differential expression analysis of genes across different age stages

To facilitate downstream analyses, we first constructed pseudo-bulk expression profiles based on donor age and brain region. Genes expressed in at least 50% of the samples were retained for analysis. To account for potential confounding effects of sex and brain region on the identification of age-related genes, we built a design matrix and applied a linear modeling framework using the limma package (v3.64.1) ^107^. For each gene, we calculated the correlation with age, identifying those that demonstrated significant relationships. Ultimately, we identified 1,814 age-related genes, each exhibiting an absolute correlation coefficient with age greater than 0.6 in at least one layer. This approach ensured a comprehensive evaluation of the factors influencing gene expression across different ages and layers.

#### Co-expression network analysis in age-related genes by hdWGCNA

We conducted a gene co-expression network analysis of spatial transcriptome data using the hdWGCNA package (v0.4.03) ^38^, focusing on 1,814 age-related genes. For each layer and stage, we implemented a bootstrapped cell aggregation procedure to construct metacell-level gene expression profiles. This involved pooling 20 bin100 segments per metacell, determined through K-nearest neighbors, using the MetacellsByGroups function. Subsequently, we computed a topological overlap matrix (TOM) to characterize the gene co-expression network and employed the Dynamic Tree Cut algorithm through the ConstructNetwork function to group genes into co-expression modules. To summarize the gene expression profiles of each co-expression module, we calculated module eigengenes (MEs) using the ModuleEigengenes function. We further refined the resulting MEs by applying Harmony to account for individual batch assignments, ensuring the robustness of our analysis.

To systematically investigate the functional specialization of cortical cell populations in gene modules across cortical layers, we implemented a multi-modal analytical approach. First, we leveraged module-specific gene signatures to comprehensively evaluate cellular heterogeneity across distinct cortical layers. Subsequently, we established a novel integrative metric, termed the Module Proportion Integrated Score (MP score), which combines transcriptional profiles of module-associated genes with cellular composition dynamics. The MP score for a given cell subtype *c* and functional gene module *α* at a specific cortical layer was defined as:

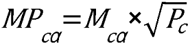

where *M_ca_* denotes the module-specific enrichment score for cell subtype *c* (calculated via gene set scoring algorithms), and *P_c_* represents the proportional abundance of cell subtype *c* within a specific cortical layer. For downstream mechanistic exploration, we implemented a rigorous selection criterion by identifying cell subtypes exhibiting statistically significant MP scores for each module. These high-confidence subtypes were prioritized for subsequent pathway analysis.

#### Identification of age-associated cell subtypes

To ensure statistical robustness, we first filtered for cell subtypes with a total count exceeding 5,000 nuclei across all samples, retaining only these for downstream analysis. For each retained cell subtype, we calculated its average relative abundance across cortical layers (L1–6) and across age stages. Relative abundance was defined as the proportion of a given cell subtype relative to the total number of cells within a specific layer and sample. To systematically evaluate the dynamic association between cell type abundance and age, linear regression models were constructed for each cell subtype and cortical layer combination, with average cell percentage as the response variable and age stage as the predictor variable. Standardized regression coefficients extracted from these models were used as key indicators to measure the direction and strength of age-related abundance changes. Based on the absolute magnitude of these regression coefficients, the top 20 cell subtypes exhibiting the most significant changes were selected and defined as age-associated cell subtypes.

#### Spatial intercellular communication analysis

We employed the StereoSiTE package (v2.2.2) ^108^ to spatially analyze cell–cell communication among the cell groups of interest in the Stereo-seq data. The analysis was based on 1,939 curated ligand–receptor interactions from the CellChatDB.human database, which classifies interactions into three categories: Secreted Signaling, ECM–Receptor, and Cell–Cell Contact. To assess communication probabilities between cell groups, we utilized the intensities_count function with a spatial distance threshold of 100 µm. Ligand–receptor pairs were filtered based on statistical significance (*P*-value < 0.05) and were required to be detected in at least four age stages. The intensity of these inferred cell–cell communications was then visualized using the ggplot2 package (v3.5.2) ^102^, enabling comprehensive representation of spatial signaling dynamics between cell types.

#### LD score regression analysis of age-related modules using LDSC

For each gene module, we employed LiftOver (https://genome.ucsc.edu/cgi-bin/hgLiftOver) ^109^ to convert genomic coordinates from the hg38 to the hg19 reference genome assembly. To compute the module-specific linkage disequilibrium (LD) scores, we generated annotation files for each gene module across all 22 autosomes, which served as input for the LD score calculation. The detailed workflow for this process can be found in the LDSC (v1.0.1) ^42,110^ documentation (https://github.com/bulik/ldsc/wiki/Cell-type-specific-analyses). Finally, we assessed the association between each gene module and the feature of interest by using the *P*-value of the regression coefficient as the statistical measure of significance.

#### Variance partition analysis of Stereo-seq and snRNA-seq data

To investigate the contribution of biological and technical factors to gene expression variability, we performed variance partition analysis on both Stereo-seq and snRNA-seq datasets. For the spatial transcriptomic data, we first grouped cell-bin data by section, cortical layer, and cell subclass, excluding groups with fewer than 10 cells. Within each group, we generated metacells by aggregating the transcriptomes of the 10 nearest cells (allowing duplication). SCTransform normalization was applied on each section, followed by random sampling of 10 metacells per group to control data size. Genes with maximum expression ≤ 5 UMIs or no variation across metacells were removed. Only genes expressed in more than 15 metacells were retained to reduce stochastic noise. Cell subclasses “SST CHODL” and “L5 ET” were remove from data to ensure our data have enough cells for each subclass. Variance partitioning was performed using the variancePartition package (v1.28.0) ^43^, based on the SCTransform-normalized expression matrix. We included the following factors in the variance model: donor, subclass, layer (L1–6), sex, cortical lobe (FL, PL, TL, OL), pathology (epilepsy, abscess, tumor, carcinoma, other), and age (≤ 12, 13–20, 21–40, 41–60, ≥ 61 years). Genes with ≥ 3% variance explained by any factor were retained for downstream analysis. For the snRNA-seq dataset, the same variance partitioning framework was applied, except that the “layer” factor was excluded due to the lack of spatial information in snRNA-seq.

To identify genes strongly associated with individual variance factors, we extracted the intersection of the top 300 genes (ranked by variance proportion attributed to each factor) from both the snRNA-seq and Stereo-seq variance partitioning results. The top 30 overlapping genes that were retained in the Stereo-seq dataset were selected for demonstration.

To assess the relative variance influence of cell subclass and cortical layer on gene expression, we defined the relative variance ratio as follows:

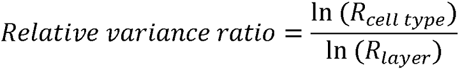

Where *R_Cell type_* denotes the proportion of gene expression variance attributed to cell type (subclass), and *R_layer_* denotes the proportion attributed to cortical layer.

#### Curation of housekeeping, drug target, and disease-related gene sets

Housekeeping genes were obtained from the Housekeeping and Reference Transcript Atlas (HRT Atlas) ^48^, specifically from the “Human housekeeping genes/transcripts and reference transcripts” download section. The file “Housekeeping genes” was used, followed by removal of duplicated entries. Brain disease-related drug target genes were extracted from the Therapeutic Target Database (TTD) ^47^. Drug targets are filtered by respective diseases which filtered by ICD-11 code to restrict diseases to brain-related diseases ^96^: 8A00-8E7Z, nervous system disease; 6A00-6E8Z, mental behavioural neurodevelopmental disorder; 7A00-7B0Z, sleep wake disorder; MB20-MB9Y, mental nervous symptom. In addition, drug targets were filtered to include only those in the “Approved” stage to ensure therapeutic relevance.

Disease categories and corresponding conditions were obtained from the DISEASES database ^121^, filtered using Disease Ontology IDs (DOIDs) ^97^. Specifically, we used the following DOIDs: 150, disease of mental health; 331, central nervous system disease; 574, peripheral nervous system disease. Diseases not mapped to any of the above DOIDs were assigned to the other disease category.

#### Sample preparation and DNA extraction for whole-genome sequencing

Genomic DNA was extracted from each brain tissue sample using the QIAamp DNA Mini Kit (Qiagen) following the manufacturer’s protocol. DNA concentration and purity were assessed using a Qubit 4.0 Fluorometer (Thermo Fisher Scientific) and NanoDrop One spectrophotometer (Thermo Fisher Scientific), respectively. Samples with an A260/A280 ratio between 1.8 and 2.0 and a minimum concentration of 50 ng/μL were selected for further processing.

#### Whole-genome sequencing library preparation and sequencing

Whole-genome sequencing libraries were prepared following the MGIEasy Whole Genome Sequencing Library Prep Kit (MGI Tech) protocol, optimized for the DNBSEQ-T7 sequencing platform. Briefly, 1 µg of genomic DNA from each sample was fragmented to an average size of 350 bp using a custom Tn5 transposase (MGI Tech) and analyzed with the Agilent 2100 Bioanalyzer (Agilent Technologies). The fragmented DNA was then end-repaired, A-tailed, and ligated to MGI-specific sequencing adapters. The adapter-ligated DNA fragments were size-selected using AMPure XP beads (Beckman Coulter, A63881) to ensure a uniform size distribution. DNA nanoballs (DNBs) were generated using the MGIEasy Circularization Kit (MGI Tech, 1000005259) according to the manufacturer’s protocol. The size-selected libraries were circularized through single-stranded DNA ligation and amplified via rolling-circle replication to form DNBs. The DNBs were then loaded onto the DNBSEQ-T7 sequencing platform (MGI Tech) flow cell. Sequencing was performed using the paired-end 150 bp (PE150) mode, generating approximately 30× coverage per sample.

#### Whole-genome sequencing quality control and variant detection

Raw data were processed using MGI’s Sequencing Analysis Software (SAS). Adapters and low-quality bases (Phred < 20) were trimmed, and reads shorter than 50 bp were discarded. Cleaned reads were aligned to human reference genome hg38 using BWA (v.0.7.17-r1188) ^111^. Variant calling was performed following the GATK (v.4) workflow ^112^. This process included detecting variants on chromosomes 1 to 22 for each individual, with the use of GenomicsDB to accelerate the calling process. The chromosome-specific gvcf files for each individual were merged, and sequencing or alignment errors were corrected through the execution of MarkDuplicates, BaseRecalibrator, and ApplyBQSR steps. Variants were filtered using VariantFiltration, removing those with mapping quality (MQ) < 40, Fisher strand bias (FS) > 30, or strand odds ratio (SOR) > 3. High-quality single nucleotide variants (SNVs) were retained using SelectVariants, and individual variants were merged using MergeVcfs to obtain the final multi-sample Variant Call Format (VCF) file.

#### Genotype quality control

After generating the final VCF file, we first used VCftools (v0.1.16) ^113^ to ensure that all alleles were consistent with the hg38 reference genome. We conducted genotype quality control using PLINK2 (v2.00a6LM) ^114^, removing SNPs with detection rate below 90% and minor allele frequency (MAF) less than 0.05 across all samples. Through this stringent screening process, we identified a total of 5,910,253 SNPs retained for downstream analysis. To address potential linkage disequilibrium (LD) among SNPs, we excluded SNPs with Hardy–Weinberg equilibrium (HWE) *P*-values less than 1×10^-6^. We further applied LD pruning using PLINK2, with a sliding window of 50 SNPs. Pairwise LD (r^2^) was computed within each window, and one SNP was removed from any pair with r^2^ > 0.95. The window was then shifted by 5 SNPs, and the process repeated iteratively. Additionally, to avoid bias from unbalanced genotype distributions, we retained only SNPs for which each genotype category was represented in at least 3 individuals. Following all filtering steps, we obtained a high-confidence dataset comprising 563,004 SNPs with reduced LD structure.

#### Genotype principal component analysis

We conducted a principal component analysis on our genotype dataset using PLINK2. In this analysis, we specifically focused on the first two principal components, which were used as covariates in subsequent eQTL association analysis to control for potential biases introduced by population structure. To identify and verify the population composition of our samples, we downloaded genotype data from 3,202 reference individuals in the 1000 Genomes Project ^122^ with annotated ancestry. We performed the same quality control steps on these reference data and our genotype data to ensure both datasets had the same loci before analysis. Subsequently, we merged the filtered genotype datasets and carried out multidimensional scaling (MDS) analysis. The resulting MDS plot showed that our samples clustered tightly with the East Asian population in the 1000 Genomes dataset, confirming the genetic ancestry of our cohort.

#### Cis-eQTL mapping on Stereo-seq data

We generated the pseudo-bulk spatial expression matrices using the metacell dataset. To ensure data quality, we included only cortical layers and cell subclasses with more than 20,000 metacells for analysis. For each cortical layer or cell subclass, we aggregated gene-level UMI counts across all metacells from the same individual to generate individual-level pseudo-bulk expression profiles. Gene expression values were then normalized using the trimmed mean of M-values (TMM) method and transformed into ln(CPM + 1) to improve cross-sample comparability. To enhance analysis robustness, we retained only genes expressed (CPM > 1) in at least 10 individuals. This filtering resulted in a high-quality pseudo-bulk expression matrix containing over 20,000 genes.

To explore the relationship between gene expression and genetic variation across spatial contexts, we utilized the TensorQTL package (v1.0.9) ^115^ to perform cis-eQTL mapping. Variants within ±1 Mb of each gene’s transcription start site (TSS) were tested, including covariates such as genotype principal components, expression principal components, and individual characteristics (sex and age). Each genotype was required to be represented in at least four individuals to ensure robustness. We applied TensorQTL’s map_nominal method separately to each expression matrix (by cortical layer and cell subclass). To identify lead SNPs for each gene, we performed 10,000 permutations using the map_cis function. Multiple testing correction was conducted using calculate_qvalues, with a false discovery rate (FDR) threshold of 0.05. Based on the resulting significant cis-eQTLs, we defined cell type-specific eQTLs as those that are significant in only one cell subclass, and layer-specific eQTLs as those that are significant in only one cortical layer.

#### Processing of PCHi-C data

We downloaded the PCHiC_peak_matrix_cutoff0.tsv file from https://osf.io/u8tzp/, which lists the CHiCAGO scores for all promoter capture Hi-C (PCHi-C) interactions with a score of ≥ 0 in at least one cell type ^67,123^. The matrix includes the genomic coordinates of baited regions (baits) and their corresponding interacting regions (”other ends”, or “oe”). Since the provided data is in the hg19 genome assembly, we used the LiftOver tool ^109^ to convert the coordinates to hg38 reference. We then filtered the dataset to retain only interactions involving the *COL27A1* promoter-interacting region (PIR), and we visualized the regulatory architecture of *COL27A1* in the context of 3D genome organization.

#### Cross-species analysis of layer-specific gene expression and cell–cell communication

For cross-species comparison, we utilized previously published macaque and mouse cortical Stereo-seq datasets ^20,21^, and selected regions homologous to the human superior frontal gyrus (SFG), middle temporal gyrus (MTG), superior parietal cortex (SPC), and primary visual cortex (V1). Homologous genes among the three species were obtained from the Ensembl BioMart database and used in subsequent analyses. Stereo-seq data from three species (bin100) were merged and normalized using the NormalizeData function in Seurat package with the “LogNormalize” method. The expression variance of homologous genes was estimated and partitioned using the variancePartition package, accounting for cortical region, cortical layer, and species as factors in the model. We identified 909 genes for which over 20% of expression variance could be attributed to species, and defined them as highly variable genes across species. To assess species-specific expression, we used the AverageExpression function to compute the average expression of these genes within each species. A gene was considered species-enriched if its average expression in one species was more than 1.5-fold higher than in the other two species. The proportions of such species-specific genes were visualized using pie charts, and Gene Ontology (GO) enrichment analysis was performed using the clusterProfiler package (v4.2.2) ^106^.

To identify layer-specific genes across species, we applied the FindAllMarkers function from the Seurat package to macaque and mouse Stereo-seq data (bin100), using an adjusted *P*-value (p_val_adj) threshold of < 0.05 and an average log_2_ fold change (avg_log2FC) > 0.1. Genes that met these criteria in more than 10% of individual sections within each brain region were retained for comparison with human GLEEs.

To assess differences in cell–cell communication across species, we used single-cell Stereo-seq data with cell type (subclass) annotation from three species. Each cell subclass was downsampled to approximately 5,000 cells to ensure balanced representation. Intercellular communication among cell subclasses was inferred using the CellChat package (v1.6.1) ^116^ across four brain regions. This process was repeated 100 times per brain region per species, with 90% of cells randomly sampled in each iteration. The resulting communication networks were aggregated for downstream comparative analysis.

## Acknowledgements

The project was supported by Lingang Laboratory (Grant No. LGL-5555, LGL-6672 and LGL-8888 to W.W.); National Natural Science Foundation of China (81870187, 81911530167 to W.W.); Shanghai Leading Talent Program of Eastern Talent Plan (LJ2023087 to W.W.); Program of Shanghai Academic Research Leader (23XD1450700 to W.W.); Major Science and Technology R&D Project of the Science and Technology Department of Jiangxi Province (20213AAG01013 to W.W.); National Science and Technology Innovation 2030 Major Program (2021ZD0204400 to L.H.); National Key R&D Program of China and by the National Science (2022YEF0203200, 2022YFA1603604 to Z.S.); Central Guidance Funds for Local Science and Technology Development (YDZX20233100001003 to H.Y.).

## Author contributions

Conceptualization, L.H., L.W., Z.S., Y.M., W.W.; Data curation, J.F., H.Y., X.Z., M.W., Yuyang Liu, X.S., Y.S.Z., J.L., Huanhuan Li, L. Li, Y.A., B.J., Y. Zhong, Q.C., Q.T., X.T., Y. Lin, R.Z., S.W., M. Chang, B.Y., M. Chen, L.M., L.Z., N.Y., H.G., Hao Li, W.Y.; Formal analysis, Z.L., Yuxuan Liu, Y.W., J.M., Y.H., Y. Zuo, J.F., Z.W., W.L., L.G., J.L.; Investigation, Chao Li, T.H., X.L., C. Liu, Y.S., L.C., L. Liu, X.X.; Methodology, Z.L., Yuxuan Liu, Y.W., J.M., Y.H., Y. Zuo, J.F., Z.W., W.L., L.G.; Project administration, L.H., L.W., Z.S., Y.M., W.W.; Supervision, L.H., L.W., Z.S., Y.M., W.W.; Visualization, Z.L., Yuxuan Liu, Y.W., J.M., Y.H., Y. Zuo, J.F., Z.W., W.L., L.G.; Writing – original draft, Z.L., Yuxuan Liu, Y.W., J.M., Y.H., Y. Zuo, J.F., Z.W.; Writing – review & editing, L.H., Z.L., L.W., Chengyu Li, H.G., Hao Li, M.P., W.-B.G., J.Y., W.Y., S.L., Z.S., Y.M., W.W.

---

**Figure S1.**
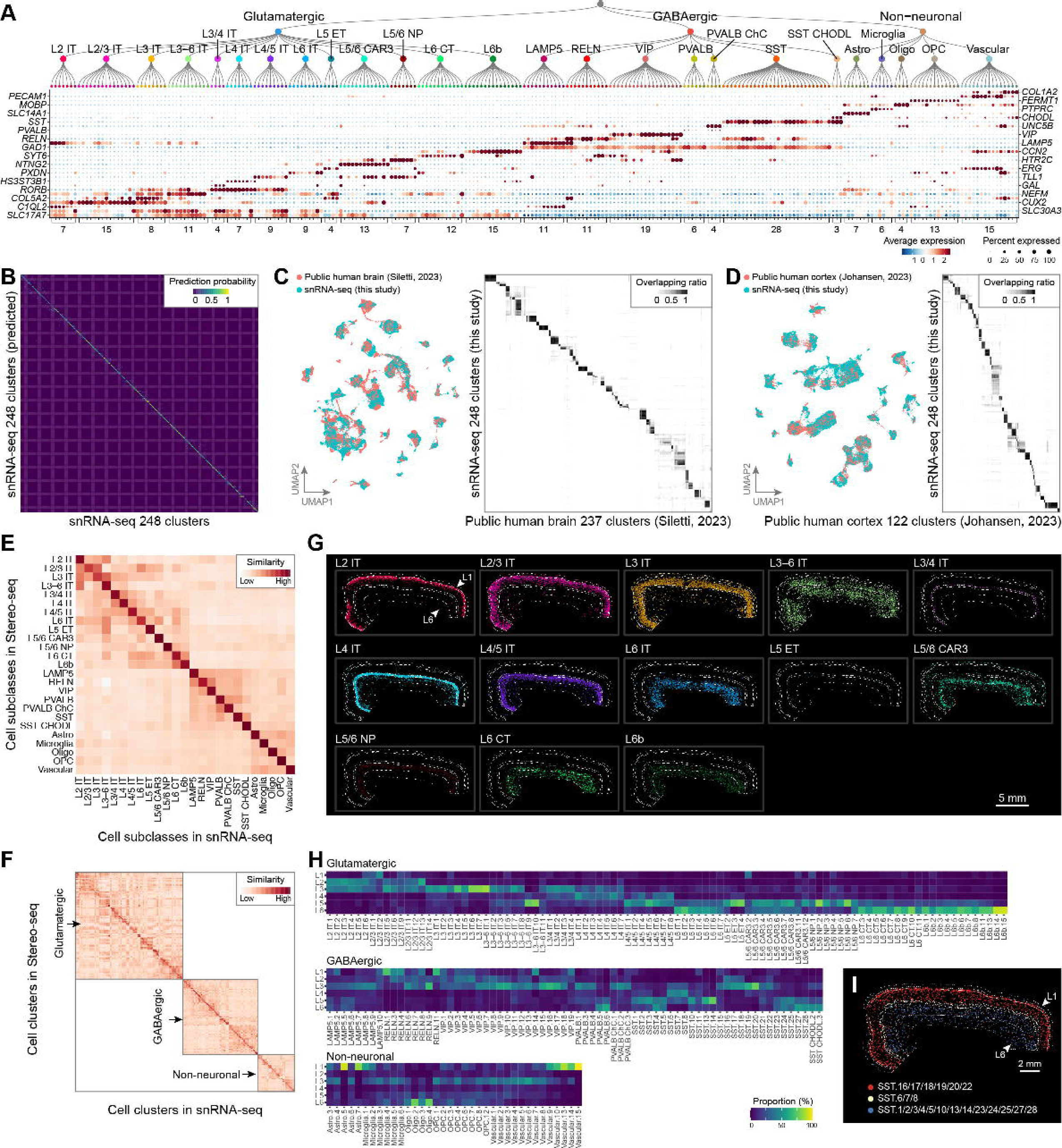
Classification and benchmark of snRNA-seq and Stereo-seq data, related to Figure 1. (**A**) Expression patterns of marker genes across 248 cell subtypes identified by snRNA-seq. Marker genes are listed on the left and right; the number of cell subtypes per subclass is indicated below. (**B**) Confusion matrix of random forest classifiers trained on the 248 snRNA-seq cell subtypes. (**C** and **D**) Comparisons of snRNA-seq cell subtypes with two published human cortex single-nucleus datasets ^8,27^. The integrated datasets visualized in UMAP space (left). Pairwise comparison of cell subtypes from integrated datasets (right). (**E** and **F**) Scaled (z-score) correlations of gene expression profiles between Stereo-seq and snRNA-seq data, across cell subclasses (**E**) and individual cell subtypes (**F**). (**G**) Spatial distribution of glutamatergic neurons subclasses on a human cortical section (T991). Scale bar, 5 mm. (**H**) Proportion of cell subtypes across cortical layers in Stereo-seq data. (**I**) Spatial distribution of layer-specific SST neurons subtypes on a human cortical section (T991). Scale bar, 2 mm.

**Figure S2.**
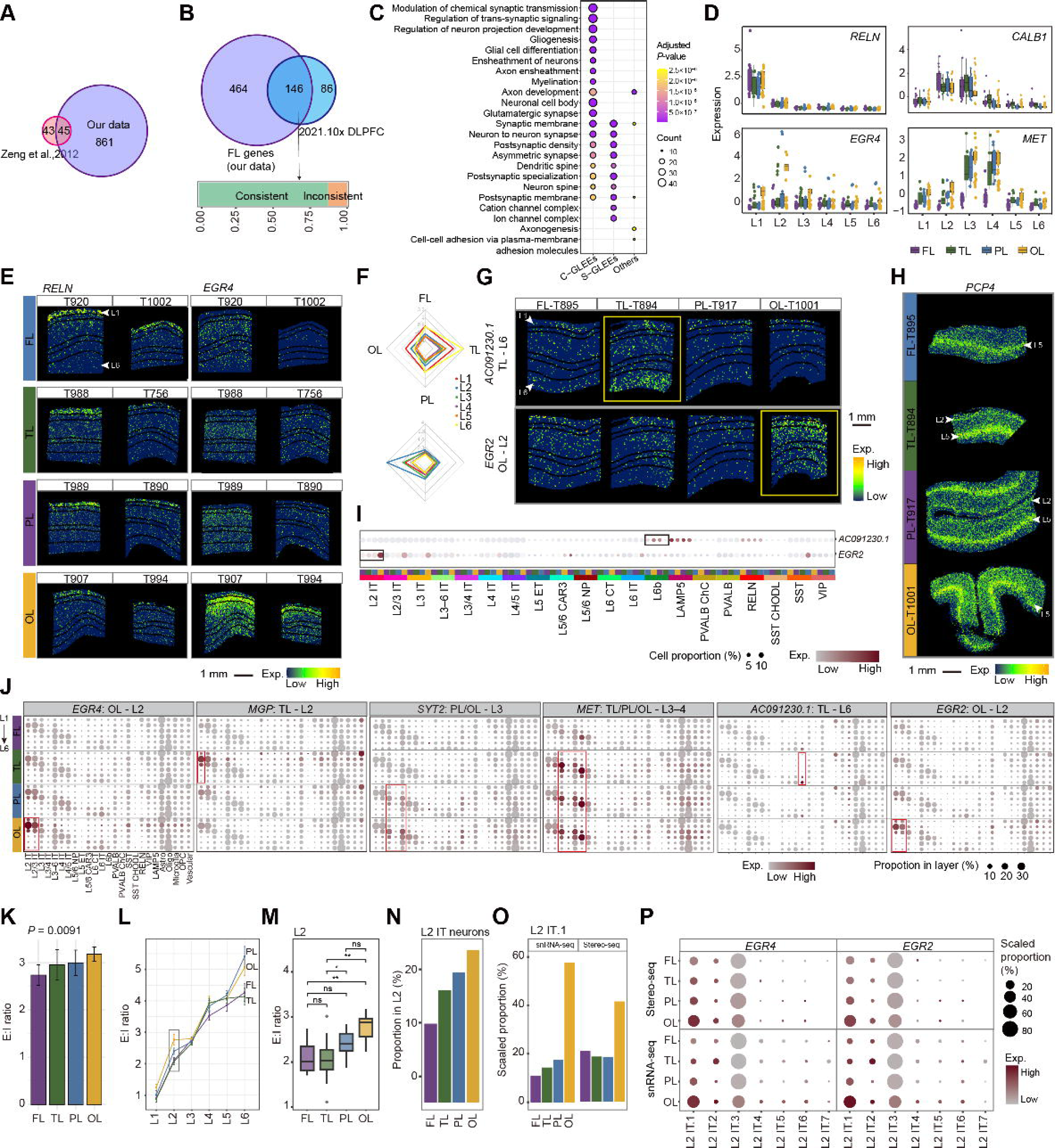
Diversity of GLEE distribution and cell composition across four cortical lobes, related to Figure 2. (**A**) Venn diagram showing the overlap between GLEEs identified in this study and those previously reported in the OL ^29^. (**B**) Venn diagram showing the consistency of GLEEs between 10x Genomics Visium data ^13^ and Stereo-seq in FL, analyzed under identical screening criteria. (**C**) Dot plot displaying the Gene Ontology (GO) enrichment results of GLEEs. (**D**) Box plot showing the expression levels of representative GLEEs from Stereo-seq sections. (**E**) Spatial expression patterns of representative GLEEs in additional Stereo-seq sections from the four cortical lobes. (**F**) Radar chart showing the laminar expression profiles of selected S-GLEEs across the four lobes. (**G**) Spatial expression pattern of the S-GLEEs in panel F. Scale bar, 1 mm. (**H**) Full-view spatial expression pattern of *PCP4* in the Stereo-seq sections across the four lobes. Scale bar, 1 mm. (**I**) Dot plots showing the expression levels of the S-GLEEs from panel F in the snRNA-seq data. (**J**) Dot plot showing the expression levels (color) of selected S-GLEEs across cell subclasses in each cortical layer and lobe. Dot size represents the proportion of each subclass, calculated as the number of cells of that subclass divided by the total number of cells in the same layer and lobe. (**K**) Bar plot showing the ratio of glutamatergic (excitatory) neurons to GABAergic (inhibitory) neurons (E:I ratio) in each cortical lobe (mean ± SE). (**L**) Line chart showing the E:I ratio across cortical layer in each lobe (mean ± SE). (**M**) Box plot comparing the E:I ratios in cortical L2 across four lobes. Statistical significance assessed using the Wilcox test. Significant levels are labeled (* for *P*-value < 0.05, ** for *P*-value < 0.01, n.s. for *P*-value ≥ 0.05). (**N**) Bar graph showing the proportion of L2 IT neurons in L2 across the four lobes. (**O**) Bar plot showing the scaled proportion of the L2 IT.1 subtype within L2 IT neurons across lobes, derived from snRNA-seq and Stereo-seq data. Scaled proportion was calculated as the number of cells in the subtype divided by the total number of cells in the corresponding subclass within each lobe, and subsequently normalized such that the sum across all lobes equals 1. (**P**) Dot plot showing the expression levels (color) of *EGR2* and *EGR4* in cell subtypes of L2 IT neurons across cortical lobes, based on snRNA-seq and Stereo-seq data. Dot size represents the scaled proportion of each subtype (the sum across cell subtypes equals 1).

**Figure S3.**
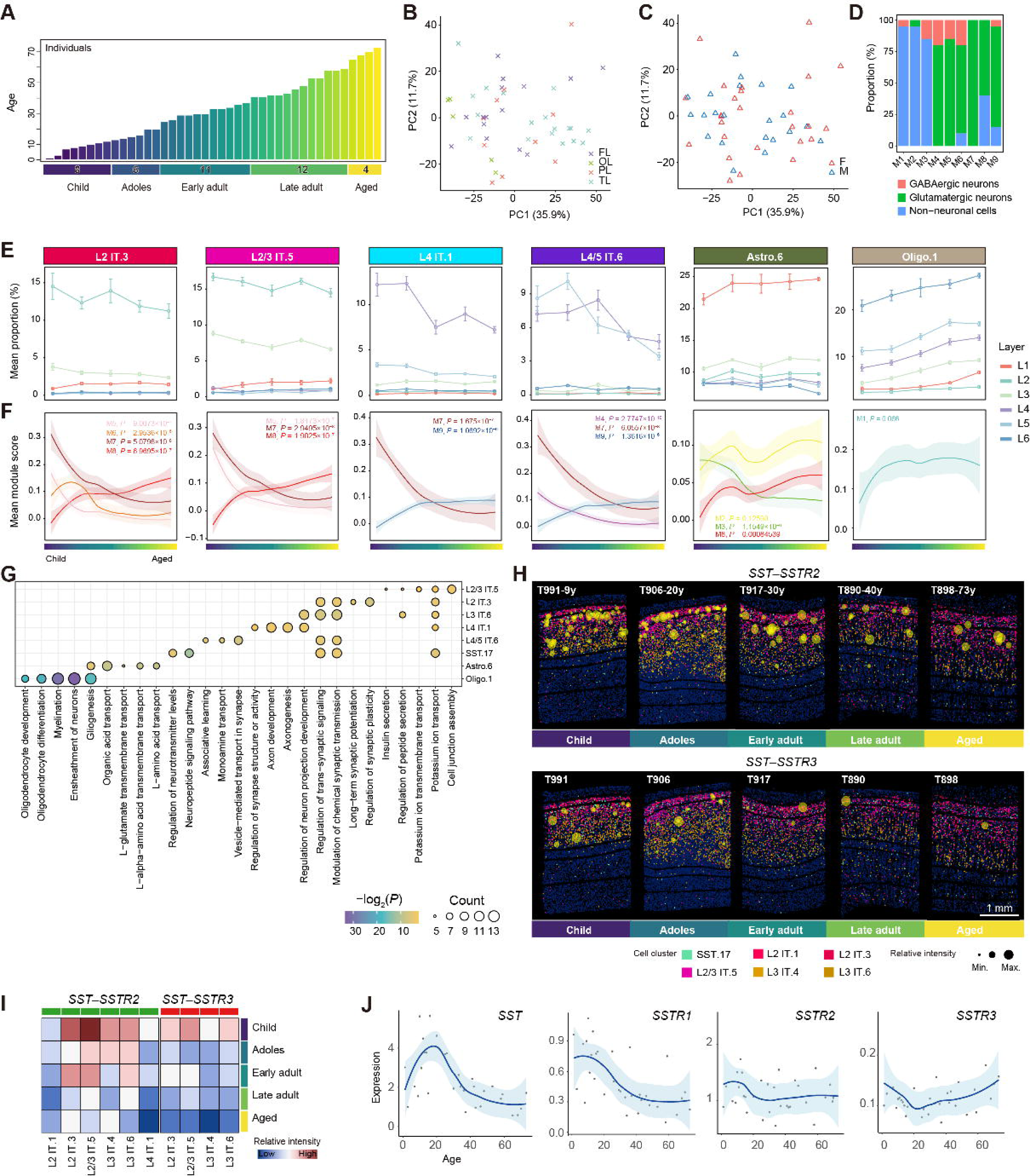
Age-related changes in cell subtypes, ARG modules, and cell–cell interactions, related to Figure 3. (**A**) Bar plot showing the age distribution of donors. (**B** and **C**) PCA plot showing the distribution of donor Stereo-seq sections colored by cortical lobes (**B**) or gender (**C**), based on the expression of all ARGs. (**D**) Bar plot showing the cell class proportion of the top 20 cell subtypes (ranked by the Module Proportion Integrated Score; **STAR Methods**) within each module. Modules were defined as neuron-associated or glia-associated if the proportion of neurons or non-neuronal subtypes exceeded 75%. (**E**) Line chart showing the proportion of age-related cell subtypes across layers and aging stages. (**F**) Line chart showing the module scores of corresponding modules related to cell subtypes in panel D across aging stages. (**G**) Dot plot showing the enriched GO terms for age-related cell subtypes in panel D, based on snRNA-seq data. (**H**) Spatial visualization showing cell–cell interaction intensity of the *SST*–*SSTR2* and *SST*–*SSTR3* ligand–receptor pairs across aging stages. Yellow circle size reflects the interaction strength. Scale bar, 1 mm. (**I**) Heatmap showing cell–cell interaction intensity of the *SST*–*SSTR2* and *SST*–*SSTR3* ligand–receptor pairs across cell subtypes and aging stages. (**J**) Line plot showing the expression levels of *SST*, *SSTR1*, *SSTR2* and *SSTR3* across aging stages.

**Figure S4.**
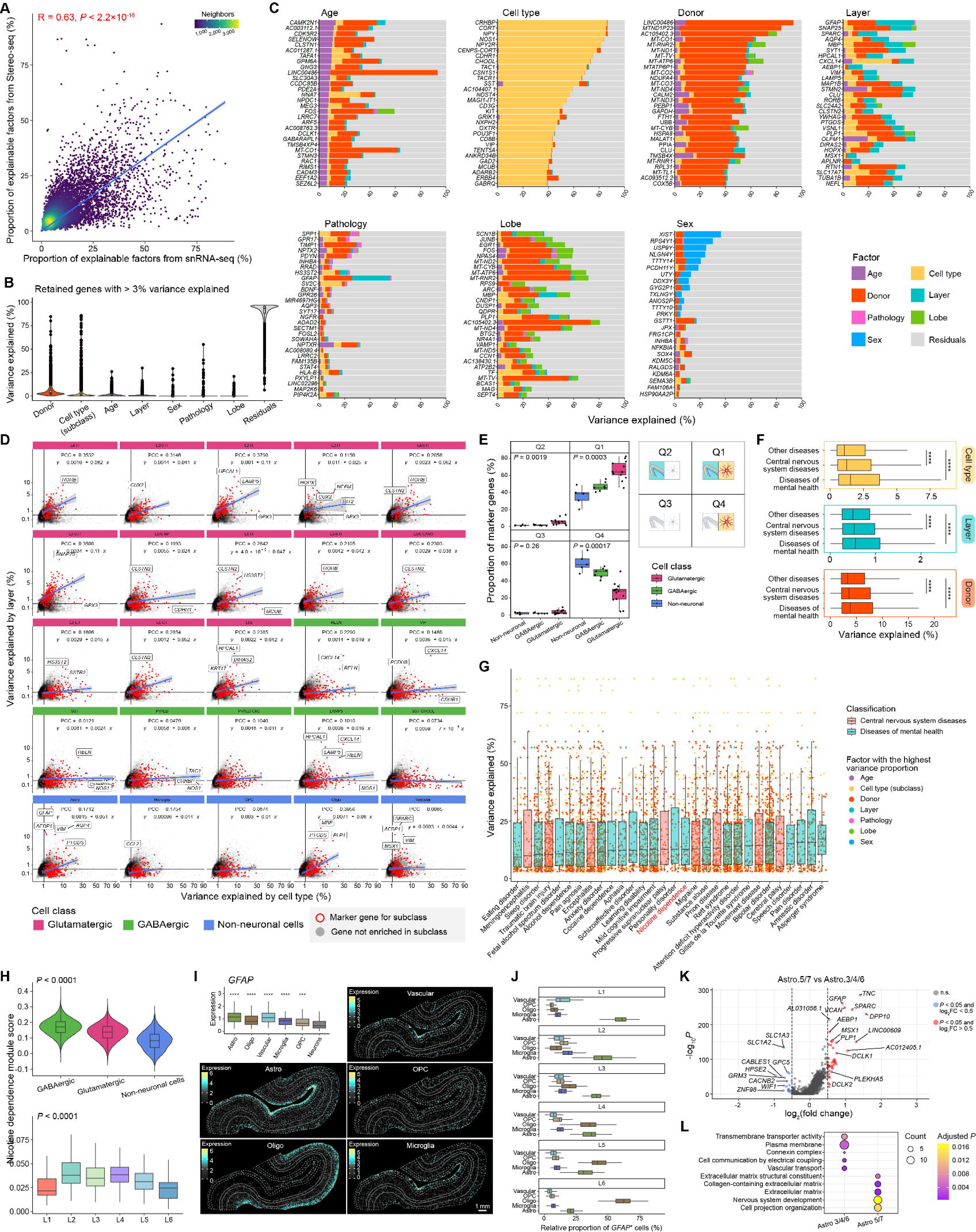
Multi-factor analysis reveals cell type and layer-related patterns of transcriptional variance, related to Figure 4. (**A**) Scatter plot showing Pearson correlation of the sum of proportions of variance across all explained factors between Stereo-seq meta-cell data and snRNA-seq data. (**B**) Faceted bar plots displaying representative genes with the highest variance (top 30, shared between Stereo-seq and snRNA-seq data) explained by specific factors. (**C**) Violin plot showing the proportion of variance explained by all annotated factors across all genes included in the analysis. Genes with > 3% variance explained were retained for downstream analysis. (**D**) Scatter plot showing variance attributed to cell type (x-axis) and cortical layer (y-axis) for all genes. Marker genes of cell subclasses (adjusted *P*-value < 0.05 and log_2_(fold change) > 1, Wilcoxon Rank Sum test with Bonferroni correction) were colored in red and other genes are colored in light grey. Linear regression lines with 95% confidence intervals are fitted for each subclass, with corresponding Pearson correlation coefficients indicated. (**E**) Comparison of the proportion of subclass-specific marker genes across the four variance-based quadrants defined in panel D. Statistical comparisons were performed using the Kruskal–Wallis test; *P*-values are labeled. (**F**) Comparison of explainable variance across disease of mental health and central nervous system diseases gene sets to other diseases (brain-irrelevant) for three major factors (donor, cell type and cortical layer). Statistical tests were carried out using Wilcoxon rank-sum test with Benjamini–Hochberg adjustment, with significant levels labeled (* for *P*-value < 0.05, ** for *P*-value < 0.01, *** for *P*-value < 0.001, **** for *P*-value < 0.0001, n.s. for *P*-value ≥ 0.05). (**G**) Box plots showing the top 30 brain-related disease gene sets ranked by median variance explained. Genes are color-coded based on the factor contributing the highest proportion of variance. (**H**) Distribution of nicotine dependence module scores across cell subclasses (top) and cortical layers (bottom), based on snRNA-seq and Stereo-seq data, respectively. Statistical comparisons were conducted using the Kruskal–Wallis test. *P*-values are shown. (**I**) Box plot and spatial visualization of *GFAP* expression in non-neuronal subclasses. Box plot comparing *GFAP* expression in non-neuronal subclasses versus neurons across Stereo-seq sections. Statistical significance was tested using the Wilcoxon rank-sum test, with significance levels as defined in panel F. Scale bar, 1 mm. (**J**) Box plot showing the relative proportion of *GFAP*^+^ cells in non-neuronal subclasses across cortical layers. The relative proportion is defined as the number of *GFAP*L cells within a given subclass divided by the total *GFAP*L non-neuronal cells in the same layer. (**K**) Volcano plot showing the differentially expressed genes (DEGs) in Astro.5/7 comparing to Astro.3/4/6. Upregulated genes (adjusted *P*-value < 0.05 and log_2_(fold change) > 0.5, Wilcoxon Rank Sum test with Benjamini–Hochberg correction) are colored in red and downregulated genes (log_2_(fold change) < -0.5) are colored in blue. (**L**) Dot plot of GO enrichment analysis based on DEGs identified in panel K.

**Figure S5.**
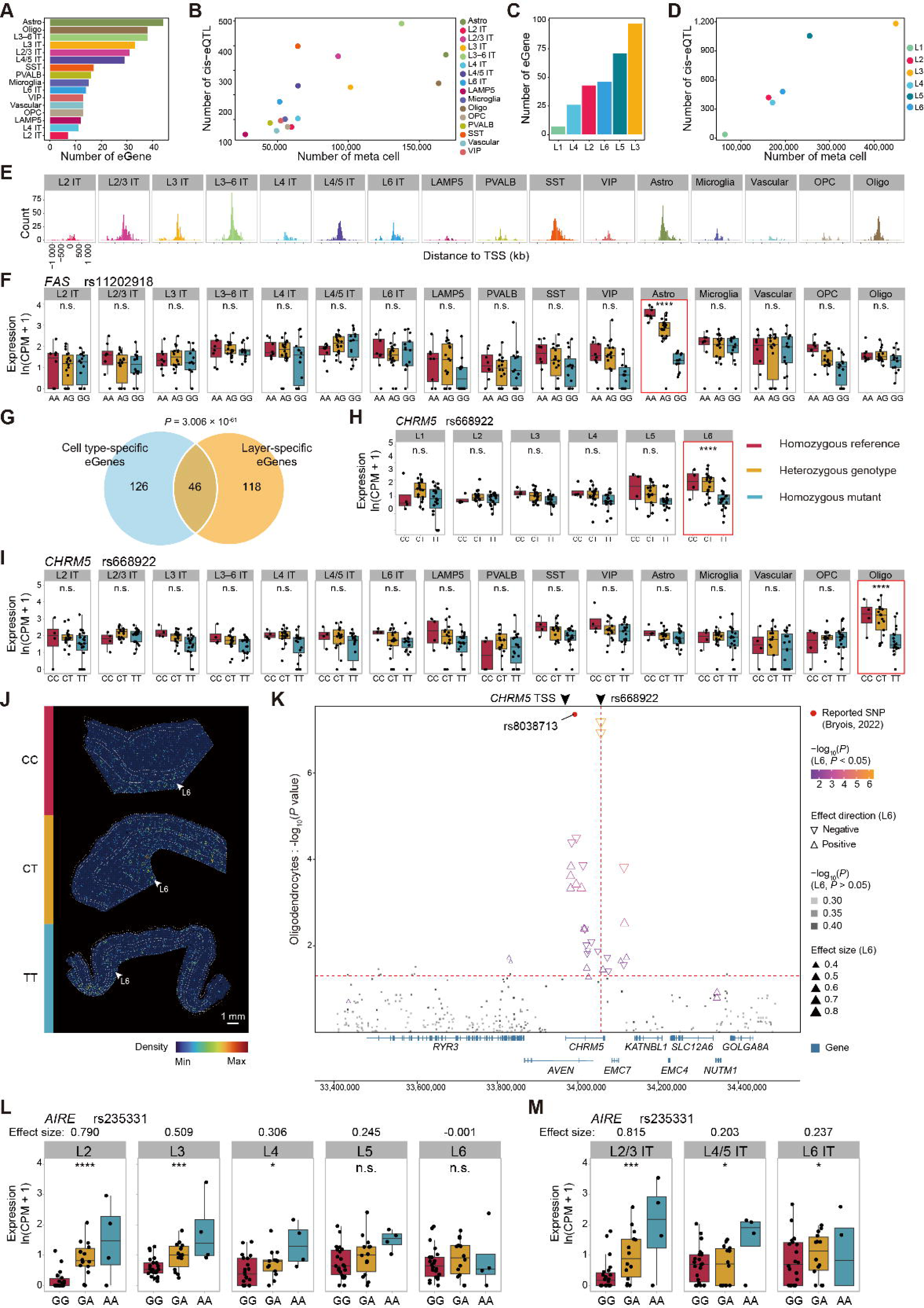
Quantitative landscape and spatial specificity of cis-eQTLs, related to Figure 5. (**A**) Number of eGenes identified across cell subclasses. (**B**) Relationship between the number of significant cis-eQTLs (FDR < 0.05) and the number of meta cells across cell subclasses. (**C**) Number of eGenes identified across cortical layers. (**D**) Relationship between the number of significant cis-eQTLs (FDR < 0.05) and the number of metacells across cortical layers. (**E**) Distribution of distances to transcription start sites (TSSs) from significant (FDR < 0.05) cis-eQTLs across cell subclasses. (**F**) Box plot showing the expression of *FAS* on various rs11202918 genotypes across cell subclasses. The significant (FDR < 0.05) cell type-specific cis-eQTL is marked in red, with *P*-value labeled (* for *P*-value < 0.05, ** for *P*-value < 0.01, *** for *P*-value < 0.001, **** for *P*-value < 0.0001, n.s. for *P*-value ≥ 0.05). (**G**) Venn diagram showing the overlap between eGenes associated with cell type-specific cis-eQTLs and those associated with layer-specific cis-eQTLs (hypergeometric test). *P*-value is shown. (**H**) Box plot showing the expression of *CHRM5* on various rs668922 genotypes across cortical layers. The significant (FDR < 0.05) layer-specific cis-eQTL is labeled by a red box, with *P*-value labeled. Significance levels as defined in panel F. (**I**) Box plot showing the expression of *CHRM5* on various rs668922 genotypes across cell subclasses. The significant (FDR < 0.05) cell type-specific cis-eQTL is marked in red, with *P*-value labeled. Significance levels as defined in panel F. (**J**) The expression density of *CHRM5* on various rs668922 genotypes in representative Stereo-seq sections (T906, T989 and T994). Highlighting bin100 where *CHRM5* expression is greater than 0, with darker colors indicating a higher local density of such expression-positive bins. (**K**) Distribution of SNPs surrounding *CHRM5* (chr15: 33,400,000-34,500,000), with a red vertical line indicating the genomic position of rs668922. A black dashed vertical line marks the transcription start site (TSS) of *CHRM5*. The y-axis displays the significance level of cis-eQTLs related to *CHRM5* in oligodendrocytes, with a red horizontal line representing the significance threshold (*P*-value = 0.05). Each triangle represents a SNP, colored by the significance levels of cis-eQTLs related to *CHRM5* in cortical L6, with size representing effect size and direction indicating the effect of regulation. (**L**) Box plot showing the expression of *AIRE* on various rs235331 genotypes across cortical layers. Effect sizes (regression slopes) and *P*-values are indicated. Significance levels as defined in panel F. (**M**) Box plot showing the expression of *AIRE* on various rs235331 genotypes across cell subclasses. Effect sizes (regression slopes) and *P*-values are indicated. Significance levels as defined in panel F.

**Figure S6.**
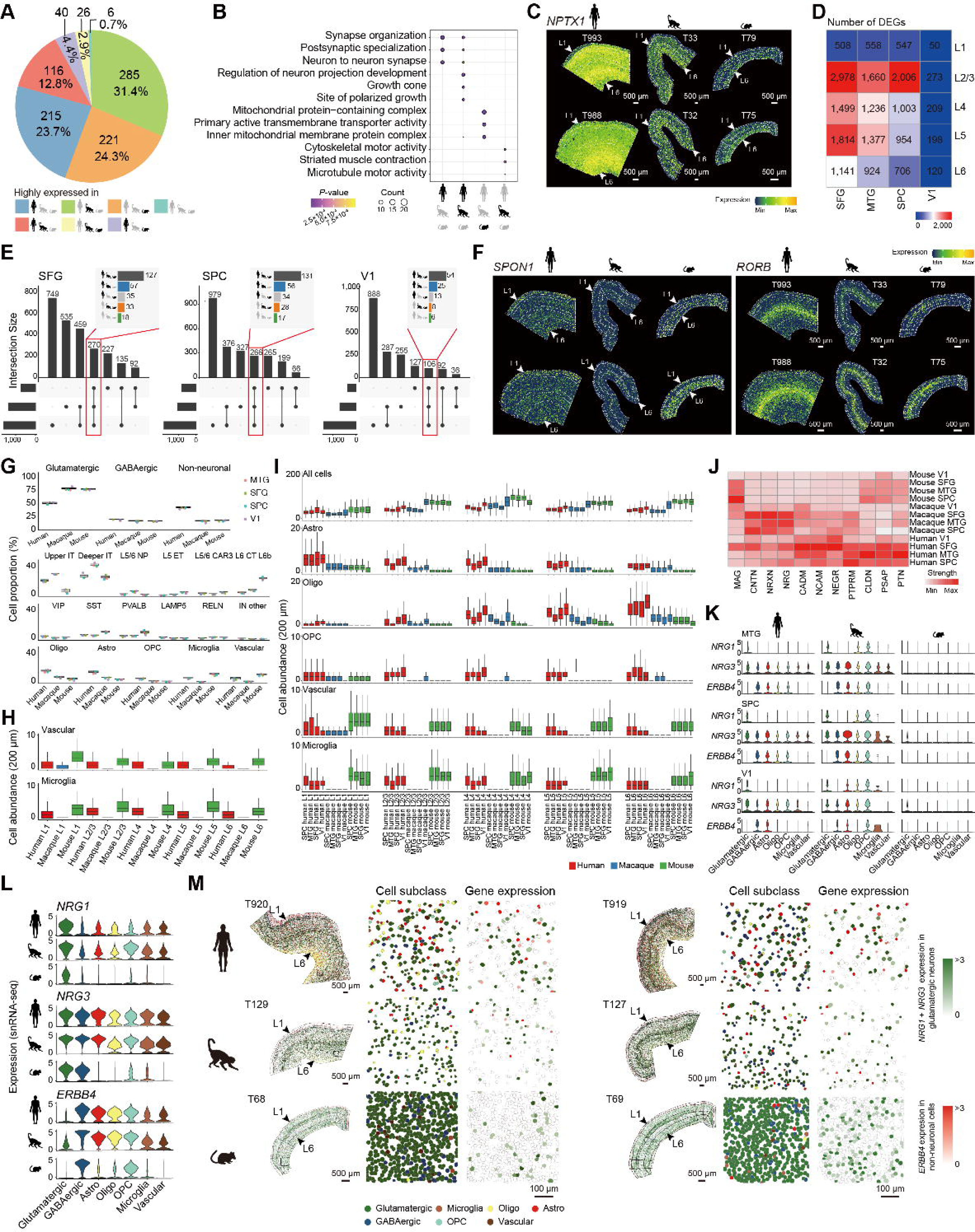
Cross-species comparison of cortical gene expression and cell composition patterns, related to Figure 6. (**A**) Pie chart showing the distribution of 909 species-associated genes (the percentage of species-related variance > 20%) based on their relative expression levels across human, macaque, and mouse. Each color represents a group of genes with preferential expression in one species, shared high expression in two species, or similar expression across all three species. (**B**) Dot plot showing the GO enrichment results of species-associated genes at various groups, including genes highly expressed in human, macaque, or mouse, as well as those shared between human and macaque. (**C**) Spatial expression patterns of *NPTX1* in the cortex of human, macaque, and mouse, as visualized in representative Stereo-seq sections. Scale bar, 500 µm. (**D**) Heatmap displaying the number of differentially expressed genes across cortical regions and layers in three species. (**E**) UpSet plot illustrating the intersection of layer-specific genes in the SFG, SPL and V1 regions across three species. The x-axis denotes intersection groups, and the y-axis indicates the number of genes per group. The consistency of layer-specific genes shared across all three species is highlighted in the bar plot, showing the number of genes with conserved or divergent laminar localization patterns. (**F**) Spatial expression patterns of *RORB* and *SPON1* in the cortex of human, macaque, and mouse, as visualized in representative Stereo-seq sections. Scale bar, 500 µm. (**G**) Box plot showing the proportions of cell classes and subclasses across four cortical regions in three species. (**H**) Box plot showing the abundance of specific cell subclasses around glutamatergic neurons across cortical layers. Cell abundance was quantified by the number of cells within a 200 µm radius. (**I**) Box plot showing the abundance of all cell types and specific cell subclasses around glutamatergic neurons across cortical regions. Cell abundance was quantified by the number of cells within a 200 µm radius. (J) Heatmap displaying communication probabilities of signaling pathways across cortical regions in three species. Communication probabilities were calculated by summing the probabilities of all ligand–receptor interactions associated with each pathway. (**K** and **L**) Violin plot illustrating the expression levels of three NRG pathway genes (*NRG1*, *NRG3*, *ERBB4*) in glutamatergic neurons, GABAergic neurons, and subclasses of non-neuronal cells in the MTG, SPL and V1 region based on Stereo-seq data (**K**). Expression profiles of these genes from snRNA-seq data are also shown (**L**). (**N**) Spatial distribution of glutamatergic neurons, GABAergic neurons, and subclasses of non-neuronal cells with highlighted expression of NRG pathway genes. The combined expression of *NRG1* and *NRG3* in glutamatergic neurons is represented with a color gradient from white to green, while *ERBB4* expression in non-neuronal cells is displayed with a gradient from white to red. Scale bar, 500 µm; zoom-in regions, 100 µm.

